# Concurrent assessment of neurometabolism and brain hemodynamics to comprehensively characterize the functional brain response to psychotropic drugs: an S-ketamine study

**DOI:** 10.1101/2024.07.23.604721

**Authors:** Daphne E Boucherie, Liesbeth Reneman, Jan Booij, Rogier Immink, Markus W Hollmann, Anouk Schrantee

## Abstract

Neuroimaging techniques are crucial for understanding pharmacological treatment effects in neuropsychiatric disorders. Here, we present a novel approach that simultaneously assesses hemodynamic and neurometabolic brain responses to psychotropic drugs using interleaved pharmacological magnetic resonance imaging (phMRI) and magnetic resonance spectroscopy (phMRS). This method was tested using a double-dose, placebo-controlled, randomized, crossover design using S-ketamine, and acquiring 7 Tesla phMRI and phMRS data to evaluate time- and dose-dependent effects in 32 healthy controls. S-ketamine elicited robust phMRI responses in the dorsal-frontal, cingulate, and insular cortices, which correlated with glutamate and opioid receptor maps and subjective dissociation scores. These hemodynamic changes were paralleled by increases in glutamate and lactate, especially at higher doses. Furthermore, accuracy in predicting received S-ketamine dose increased when combining both techniques. Here, we show for the first time that concurrent phMRI and phMRS assessments provide important complementary insights into the functional brain response to pharmacological interventions.

## Introduction

Imaging biomarkers may play an essential role in evaluating whether a drug is effectively modulating the intended neural pathways or neurotransmitter systems associated with neuropsychiatric disorders. The development of pharmacological treatments in neuropsychiatry faces a significant obstacle due to the absence of such precise biomarkers. Therefore, imaging methods that can non-invasively assess the brain’s response to medication are urgently needed^1^.

An established method showing potential in this regard is pharmacological magnetic resonance imaging (phMRI), due to its non-invasive nature, wide availability, good spatial resolution and whole brain coverage. PhMRI measures hemodynamic changes in response to pharmacological stimulation of a target neurotransmitter system. In short, after oral or intravenous administration of a drug, it binds to its specific target, thereby (de)activating the neurons on which the respective receptor or transporter is located. Changes in neuronal activity then induce a cascade of neurotransmitter and metabolite release, accompanied by an increase in local cerebral blood flow (CBF), volume (CBV) and oxygenation through an intricate process called ‘neurovascular coupling’^2^. These hemodynamic changes can subsequently be measured with MRI MRI techniques, e.g. blood-oxygen-level-dependent (BOLD) functional MRI (fMRI). As such, phMRI can provide an insight into neurotransmitter function^3–5^, and is ideally suited for drug discovery studies, as well as treatment monitoring and prediction.

Despite its considerable promise, interpreting phMRI data is challenging due to multiple complex factors influencing the phMRI signal. As phMRI relies on hemodynamic signals, non-neuronal processes, such as direct effects of psychotropic drugs on the vasculature, can affect the phMRI signal. To more directly assess the effects of medication at the neurometabolic level, proton Magnetic Resonance Spectroscopy (^1^H-MRS) can be employed, which provides information about metabolic processes occurring within tissues. Specifically, functional MRS (fMRS), in which dynamic scans are acquired at short time intervals of several seconds, allows for the assessment of acute changes in neurometabolite levels in response to neuronal activation^6–8^.

Here, we introduce a novel approach called pharmacological MRS (phMRS), which applies fMRS in the context of psychopharmacology to directly measure neurometabolite changes over time, in response to a psychotropic drug challenge. PhMRS can be used to detect changes in metabolites like glutamate, lactate, aspartate, and glucose. Glutamate largely governs the hemodynamic response to increased neuronal activity, and its turnover is tightly linked to brain energetics^9^, whereas lactate, aspartate, and glucose play important roles in the neurometabolic cycle^10^.

We aimed to gain a more comprehensive understanding of the functional brain response to medication, by simultaneously assessing neurometabolic and hemodynamic contributions. Therefore, we developed and evaluated an innovative interleaved phMRI-phMRS protocol, which concurrently measures hemodynamic and neurometabolic changes in response to drug administration (Fig. 1A). We tested this approach using a double-dose, placebo-controlled, randomized crossover design with S-ketamine as the pharmacological agent. S-ketamine was selected due to its previously demonstrated efficacy in phMRI paradigms, its clinical relevance as a treatment for refractory depression^11^, its established use as a pharmacological model for psychosis^12^, and its thoroughly characterized subjective effects profile. These factors make it an ideal candidate for testing our innovative imaging method. We assessed time- and dose-dependent effects of S-ketamine on the hemodynamic response and neurometabolite dynamics, and assessed their association with subjective effects. We also examined which receptor targets underlie these effects.

Consistent with, and extending prior work^13,14^, we show a strong rapid and a slower, more sustained phMRI response across the dorsal-frontal, cingulate and insular cortices in both low and high S-ketamine doses compared to placebo, which was associated with subjective dissociation. This phMRI response pattern was significantly associated with the receptor distribution of glutamate and opioid receptors. We found that these hemodynamic changes were paralleled by increases in glutamate and lactate in the anterior cingulate cortex (ACC) for the high dose compared to placebo. Finally, we provide compelling evidence that phMRI and phMRS offer complementary information, as they independently contributed to predicting received S-ketamine dose.

**Fig. 1.**
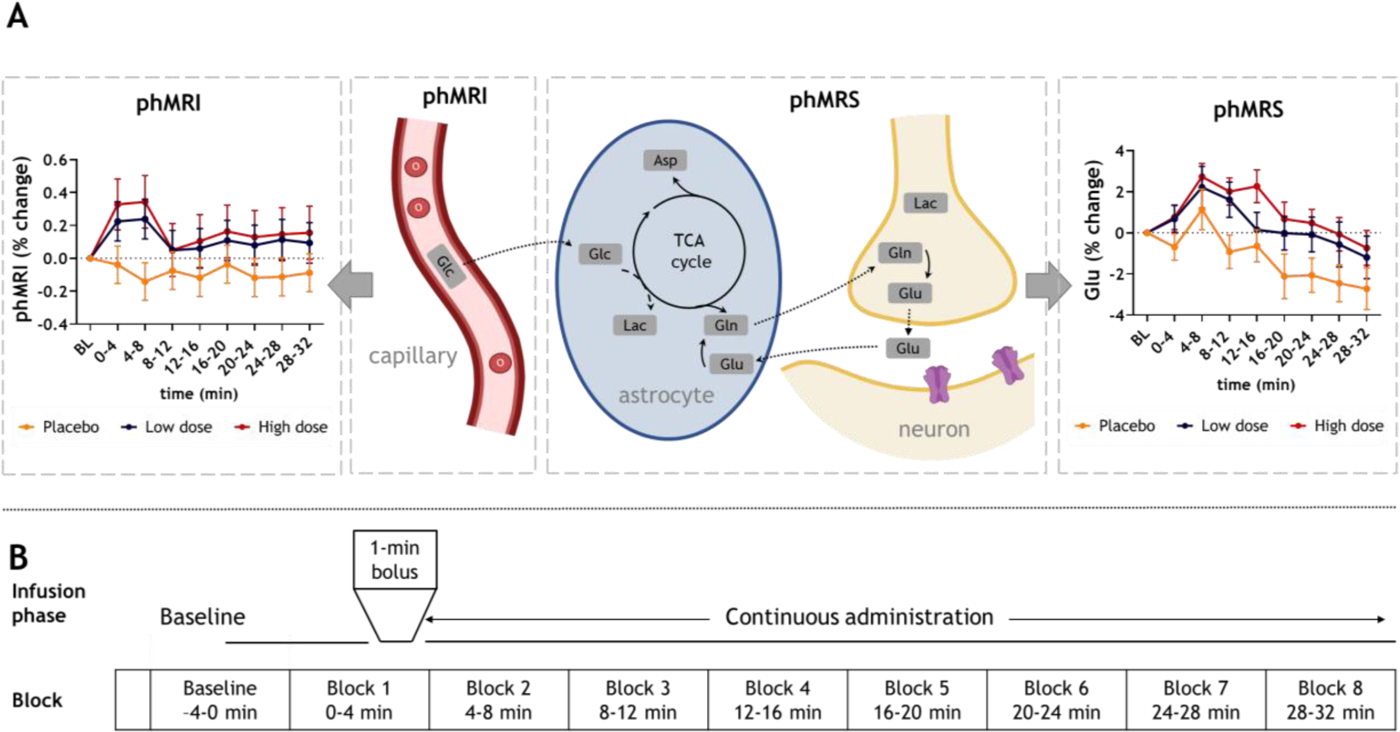
phMRI/phMRS results and acquisition scheme. **A.** *Left* phMRI: percent change in BOLD signal from baseline, extracted from the anterior cingulate cortex (ACC) MRS voxel. *Middle*: Schematic overview of complementary information provided by phMRI and phMRS. *Right*: phMRS: percent change in glutamate (Glu) from baseline is visualized per condition. All data is shown as mean+SD. **B.** Schematic representation of phMRI-phMRS sessions and allocation of blocks for analysis. After a 4-minute baseline acquisition (two minutes of data at the start was discarded), S-ketamine or placebo was administered and eight 4-minute infusion blocks were acquired. phMRI: pharmacological MRI; BL: baseline; BOLD: blood-oxygen-level-dependent; min: minute; MRS: magnetic resonance spectroscopy.

## Results

To characterize the S-ketamine functional brain response, we set-up a double-dose, placebo-controlled, randomized, crossover study. We included 32 participants that were invited for three visits, during which they received one of two sub-anaesthetic doses of S-ketamine (low dose: 0.11 mg/kg+0.1 mg/kg/h, total dose≈0.17 mg/kg; high dose: 0.22 mg/kg+0.2 mg/kg/h, total dose≈0.33 mg/kg) or placebo (NaCl 0.9%) intravenously, in a counterbalanced order. Twenty-six participants completed all sessions and were included in dose-dependent analyses, and N=27 placebo, N=26 low dose, N=29 high dose sessions were included for time-dependent analyses. Participants underwent 7 Tesla (7T) MRI scanning with a novel integrated phMRI-phMRS protocol, employing interleaved acquisition of whole-brain voxel-wise 3D-EPI fMRI data and single-voxel sLASER fMRS data in the dorsal ACC. Data were continuously acquired across the session (Fig. 1B) and, after an baseline period, S-ketamine or placebo was administered using a bolus followed by continuous administration. Blood plasma levels were obtained after the administration period. We observed no differences for blood-plasma concentrations of S-ketamine between the low and high dose, whereas we did observe significant differences for its main metabolite, nor-S-ketamine (V=109; p=0.003). This may be expected, as S-ketamine rapidly attains maximum plasma concentrations^15^, has a high clearance rate, and short elimination half-life^16^. For both phMRI and phMRS analyses, we report dose-dependent differences below. All time-dependent changes are described in the Supplementary Results.

### S-ketamine-induced phMRI response

To assess the effects of S-ketamine on the phMRI response we employed a pseudo-block design within a general linear model (GLM) for single-subject first-level analysis^13,14^. We divided the total post-baseline acquisition time into eight blocks of approximately four minutes and contrasted each administration block to baseline, analyzed using repeated-measures analyses.

At the voxel-wise level, both sub-anesthetic S-ketamine doses induced significant increases in the BOLD signals, compared to placebo, across multiple time points (Fig. 2). The spatial pattern of increased BOLD responses covers the medial frontal, (para)cingulate, and insular areas. While the phMRI response appeared less widespread and less strong for the low compared to the high dose, these differences did not reach statistical significance, possibly implying a ceiling effect of the BOLD response for the high dose condition. Additional maps and cluster information can be found in Supplementary Fig. S1 and Supplementary Tables S1-S4.

**Fig. 2.**
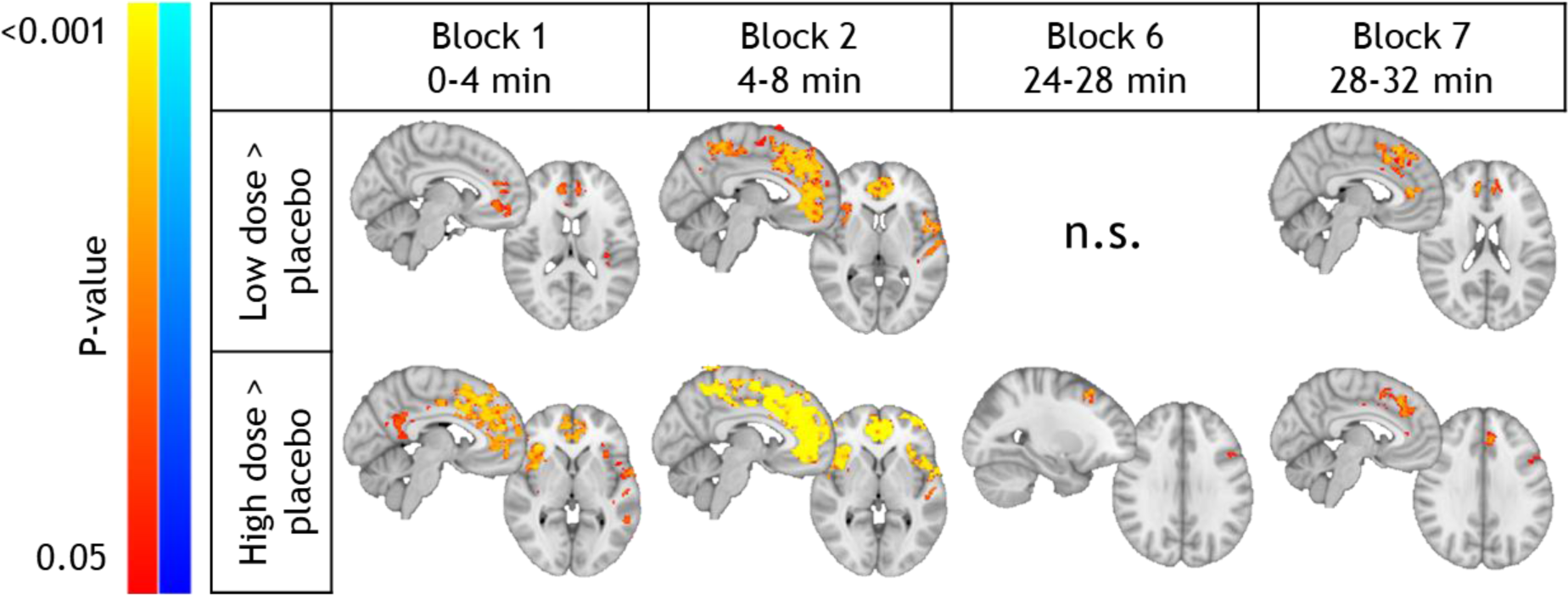
phMRI results: S-ketamine-induced effects on voxel-wise BOLD response. Clusters showing significantly higher BOLD changes (from baseline) compared to placebo for the low S-ketamine dose (top row) and high S-ketamine dose (bottom row). No significant voxel-wise differences were found for blocks 3, 4, 5 and 8. BOLD: blood-oxygen level dependent; phMRI: pharmacological magnetic resonance imaging; min: minutes.

For comparison with the fMRS data, we extracted individual-level contrast of parameter estimate (COPE) values from the location of the ACC MRS voxel and analyzed the time- and dose-dependent effects using linear mixed models. In line with the voxel-wise analysis, both S-ketamine doses elicited larger changes in BOLD responses compared to placebo (time x dose interaction; F(14,602)=1.87; p=0.027; Fig. 1A). Post-hoc comparisons showed this was the case for all infusion blocks, with the exception of block 3 (12-16 minutes post-administration). In the low S-ketamine dose, the change in BOLD in the first eight minutes was significantly higher compared to 12-20 minutes post-administration. In the high dose, the increase in BOLD in the first eight minutes was significantly larger compared to all other timepoints (Supplementary Table S5). Sensitivity analyses indicated that these results were robust when testing covariates, such as change in heart rate (Supplementary Table S6), mean motion (framewise displacement (FD), Supplementary Table S7), and order of S-ketamine administration (Supplementary Table S8). While heart rate increased during the high S-ketamine dose compared to both low dose and placebo (time*dose interaction effect, F(14,595)=2.51; p=0.002; *post-hoc*: high>placebo 8-24 minutes and high>low 12-28 minutes post-administration), this did not affect the phMRI outcomes (Supplementary Table S9).

Prior research has shown that gamma-variate models (or ‘signal model’^13,17^) are more sensitive to capture the hemodynamic response to S-ketamine compared to pseudo-block designs. Despite this, we opted for the pseudo-block design to maintain consistency across both phMRI and phMRS analyses and to characterize changes at numerous independent time points. Moreover, while we have knowledge of the hemodynamic response function, the response function of neurometabolism is less established^7,8,18^. Nonetheless, we additionally analyzed the data using a gamma-variate function (t_max_=240s and b=0.01; as per De Simoni et al.^17^; McMillan et al.^13^), and replicated the strong phMRI responses (Supplementary Fig. S2, Supplementary Tables S10-11).

While we employed the placebo condition as a control to compare against S-ketamine doses, we did observe BOLD signal fluctuations throughout the imaging session, despite regressing out cerebrospinal fluid (CSF) and white matter (WM) signals to account for physiological noise. At the acquisition, we detected diminished BOLD signals across diffuse clusters, spanning temporo-parietal regions, the paracingulate gyrus, and precuneus (Supplementary Fig. S1; Supplementary Table S1). These fluctuations might signify non-linear drifts unaccounted for by our nuisance regression or could potentially reflect changes in the participant’s cognitive state, such as a shift in attention during the session or feelings related to the absence of expected subjective effects^19^. Specifically, after placebo administration, participants may experience a release of uncertainty regarding whether ketamine was given. We therefore conducted exploratory analyses to investigate differences between participants with prior ketamine experience and those without and found that BOLD signals diminished significantly more in subjects with prior ketamine experience (N=14) compared to those without (N=13) at 4-8 minutes and 24-28 minutes post-placebo administration (Supplementary Fig. S3, Supplementary Table S12).

### S-ketamine increases glutamate levels in the ACC

Then, we investigated to what extent the hemodynamic changes induced by S-ketamine are accompanied by changes in neurometabolism. A 9 mL voxel was positioned in the dorsal ACC (Fig. 3A), motivated by prior static MRS studies hinting at ketamine’s effect on glutamate resonances in this brain region^20–22^. To quantify ketamine-induced changes, we employed spectral-temporal fitting within the dynamic MRS tool in FSL-MRS^23^. This is a novel fitting approach that was recently developed to increase the effective signal-to-noise ratio (SNR) of fMRS by analyzing all transients simultaneously. We utilize a design matrix to model the temporal response of the *concentration* of the neurometabolites, using the same pseudo-block design as for the phMRI analysis (Supplementary Fig. S4). This matrix, combined with linear combination modeling of the spectral response at each timepoint, was fitted to the entire dataset concurrently using a least-squares method^24^. While our primary analysis centered on glutamate changes, we also examined other neurometabolites demonstrating sufficient spectral quality (see Methods). As decreases in aspartate and glucose, and increases in lactate have repeatedly been observed in previous fMRS investigations due to their role in the neurometabolic cycle^10^, these neurometabolites are discussed in more detail. Additionally ascorbate, aspartate, glutathione (GSH), inositol (Ins), phosphoethanolamine (PE), scyllo-inositol, total N-acetylaspartate (tNAA), total creatine (tCr) and total Choline (tCh) are reported in the Supplementary Results. Unfortunately, gamma-aminobutyric acid (GABA) and glutamine (Gln) could not be estimated with sufficient certainty and were therefore not evaluated.

**Fig. 3.**
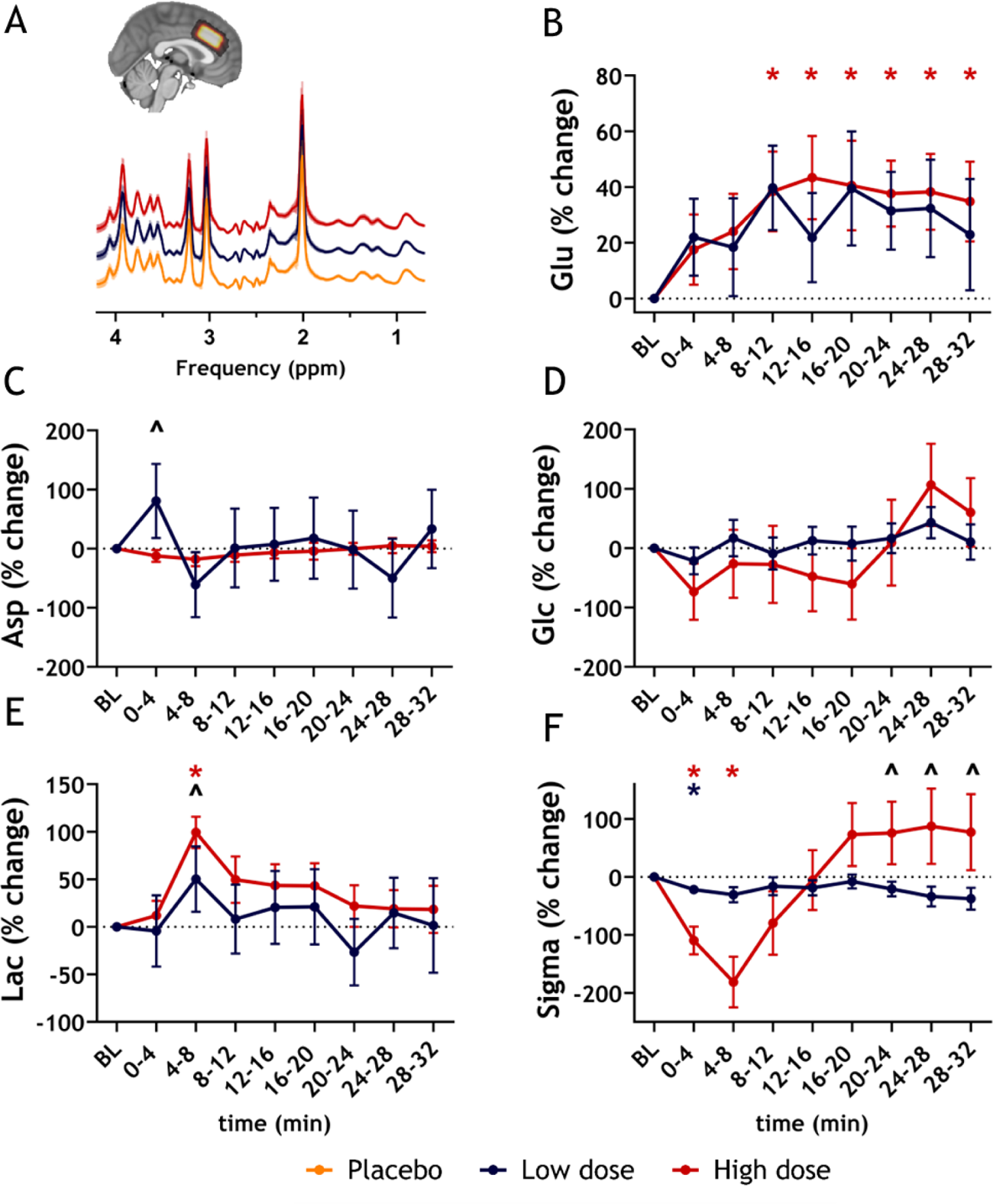
phMRS results: S-ketamine-induced changes in neurometabolite levels. **A.** Mean spectra per condition across participants (mean + standard deviation). The insert shows the overlap of ACC MRS voxel across participants (lighter is more overlap) on a 1mm MNI brain. Percentage change + standard deviation from placebo per block for **B.** glutamate (Glu) **C.** aspartate (Asp) **D.** glucose+tau (Glc) **E.** lactate (Lac) **F.** linewidth (Sigma). Differences between low dose and placebo conditions are illustrated with a blue asterisk; significant differences between high dose and placebo conditions are depicted with a red asterisk; significant differences between low and high dose conditions are depicted using a black circumflex. ACC: anterior cingulate cortex; BL: baseline; min: minute; MNI: Montreal Neurological Institute; MRS: magnetic resonance spectroscopy; ppm: parts per million.

Our results demonstrated that only the high dose of S-ketamine induced significantly higher Glu change (from baseline) compared to placebo from eight minutes after administration until the end of the acquisition (block 3-8: p_FDR_<0.05). The low dose increased Glu compared to placebo 8-12 minutes post-administration, but this did not survive multiple-comparison correction (block 3: p_FDR_=0.06). We observed no significant differences between the low and high doses after multiple-comparison correction (Fig. 1B and 3B, Supplementary Tables S13-14).

Unexpectedly, a significantly higher increase in Asp was observed for the low compared to the high dose in the first block (p_FDR_=0.02) (see Fig. 3C). No significant differences between placebo and S-ketamine conditions in Glc were found (Fig. 3D). Contrastingly, Lac increased significantly more from baseline for the high S-ketamine dose in 4-8 minutes post-administration compared to the low dose (p_FDR_=0.02) and placebo (p_FDR_<0.0001) (Fig. 3E).

Importantly, our dynamic MRS models take into account changes in the BOLD signal that lead to line narrowing in MR spectra due to alterations in T2*^10,25^. Therefore, we included a variable *linewidth* in our models (denoted as sigma, the Gaussian linewidth parameter), modeled using same design matrix as the *concentration* values with addition of a drift regressor (to capture potential effects of drift in BOLD reflected in linewidth, see Supplementary Fig. S4B). We observed that sigma mirrored the patterns observed in the phMRI analysis of the ACC voxel, namely significant decreases from baseline (reflecting increased BOLD signals) in the initial blocks after S-ketamine administration (Fig. 3F). Sigma decreased significantly in the high dose compared to the low dose (block 1-2: p_FDR_<0.05) and placebo (block1-2: p_FDR_<0.001). In the low dose, sigma also decreased compared to placebo (block 1: p_FDR_=0.048). In block 6-8, sigma increased in the high compared to the low dose (p_FDR_<0.05). (Supplementary Tables S13-14).

### phMRI and phMRS provide complementary information in distinguishing S-ketamine dose and placebo condition

Next, we examined to what extent phMRI and phMRS provide complementary information in assessing the functional brain response to S-ketamine. Therefore, we tested whether the received S-ketamine dose (condition) could be best predicted based on i) the phMRI response (Supplementary Table S15), ii) the phMRS response (Supplementary Table S16), or iii) a combination of both. We used Bayesian multilevel modeling to estimate the best-fitting models, with condition as a categorical outcome and individual-level phMRI and phMRS ACC COPE values as predictors. In the combined model, both phMRI and phMRS (Glu and Lac) predictors contributed to the best fit (Supplementary Table S16). Based on the Leave-One-Out Cross-Validation Information Criterion (LOO-IC) and the Widely Applicable Information Criterion (WAIC), the combined phMRI/phMRS model outperformed separate phMRI and phMRS models in predicting the received condition. The final model indicates that higher phMRI and higher Glu COPE values were associated with an increased probability of S-ketamine conditions compared to placebo. Additionally, Lac COPE values significantly increased the likelihood for the high S-ketamine compared to placebo. Predictors that distinguished between low and high S-ketamine doses were phMRI and Lac COPE values, as indicated by the difference in posterior distributions for these predictors between the low and high dose conditions (see Supplementary Table S17 for coefficients and difference scores).

### Dissociative experiences are associated with functional brain response to ketamine

We assessed subjective effects using the validated Clinician-Administered Dissociative States Scale (CADSS). We observed that both S-ketamine doses induced a dissociative state in participants, with significantly stronger dissociation for the high compared to the low dose (χ^2^(2)= 41.6; p<0.0001; *post-hoc* placebo vs low: Z=300; p<0.0001; placebo vs high: Z=351; p<0.0001; low vs high: Z=278.5; p=0.009) (Fig. 4A). Next, we examined whether the changes in the phMRI and phMRS responses were associated with dissociation. We associated total CADSS scores with the phMRI COPEs extracted from the ACC voxel, and the Glu COPEs using repeated-measures correlations. This revealed a significant and positive correlation between phMRI COPEs and total CADSS score in all administration blocks (Fig. 4B, Supplementary Table S17). The observed associations between Glu COPE and CADSS during administration blocks 1, 2, 3, and 6 did not survive correction for multiple comparisons (Fig. 4C, Supplementary Table S18).

**Fig. 4.**
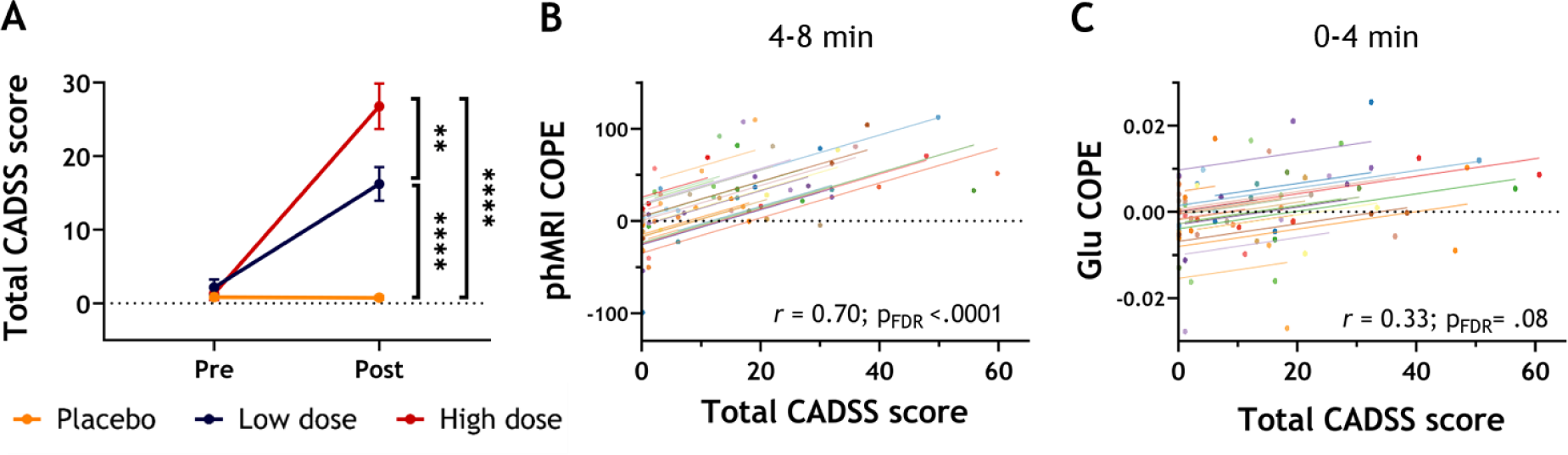
Dissociation and its association with neuroimaging measures. **A.** Total dissociation scores, as measured with the CADSS, before and after administration of placebo or S-ketamine. A dose-dependent increase in dissociation scores after administration was observed, with the high dose eliciting the highest dissociation in comparison to both low dose and placebo. **p<.01 **** p<.0001 **B.** Repeated-measures association between phMRI COPE values and total CADSS scores at 4-8 minutes post-administration. **C.** Repeated-measures association between glutamate (Glu) COPE values and total CADSS scores at 0-4 minutes post-administration. CADSS: Clinician-Administered Dissociative States Scale; COPE: contrast of parameter estimate; phMRI: pharmacological magnetic resonance imaging.

### Ketamine phMRI response is associated with glutamatergic and opioid receptor distributions

To shed light on the receptors implicated in the functional brain response to ketamine, we conducted exploratory analyses to determine the association between the phMRI response maps (placebo vs S-ketamine contrast) and neurotransmitter receptor distributions (obtained from publicly available positron emission tomography (PET) maps). Based on the results of our phMRI analysis, we investigated both the observed rapid (4-8 minutes post-administration - block 2) and the slower, or more sustained, response to S-ketamine (16-20 minutes post-administration - block 5). We tested the associations with receptor distribution maps that ketamine has been shown to target: the glutamate metabotropic receptor 5 (Glu MR5), GABA_A_ receptor, serotonin transporter (5-HTT), serotonin 2A receptor (5-HT2A), acetylcholine transporter (VAChT), metabotropic acetylcholine receptor 1 (mAChR M1), dopamine transporter (DAT), dopamine D2 receptor (D2), norepinephrine transporter (NET), and μ and κ opioid receptors^26^. These associations were evaluated using a spin-test^27,28^, a statistical method employed to compute null models by randomly rotating data, allowing for the assessment of statistical significance between two spatial maps^29^. For the rapid response, we observed a consistent association with the Glu MR5 and κ pioid receptor for both S-ketamine doses. For the high dose, the rapid response was additionally associated with the μ opioid receptor map. The sustained response was only associated with the Glu MR5 map for both S-ketamine conditions (Fig. 5 and Supplementary Table S19).

**Fig. 5.**
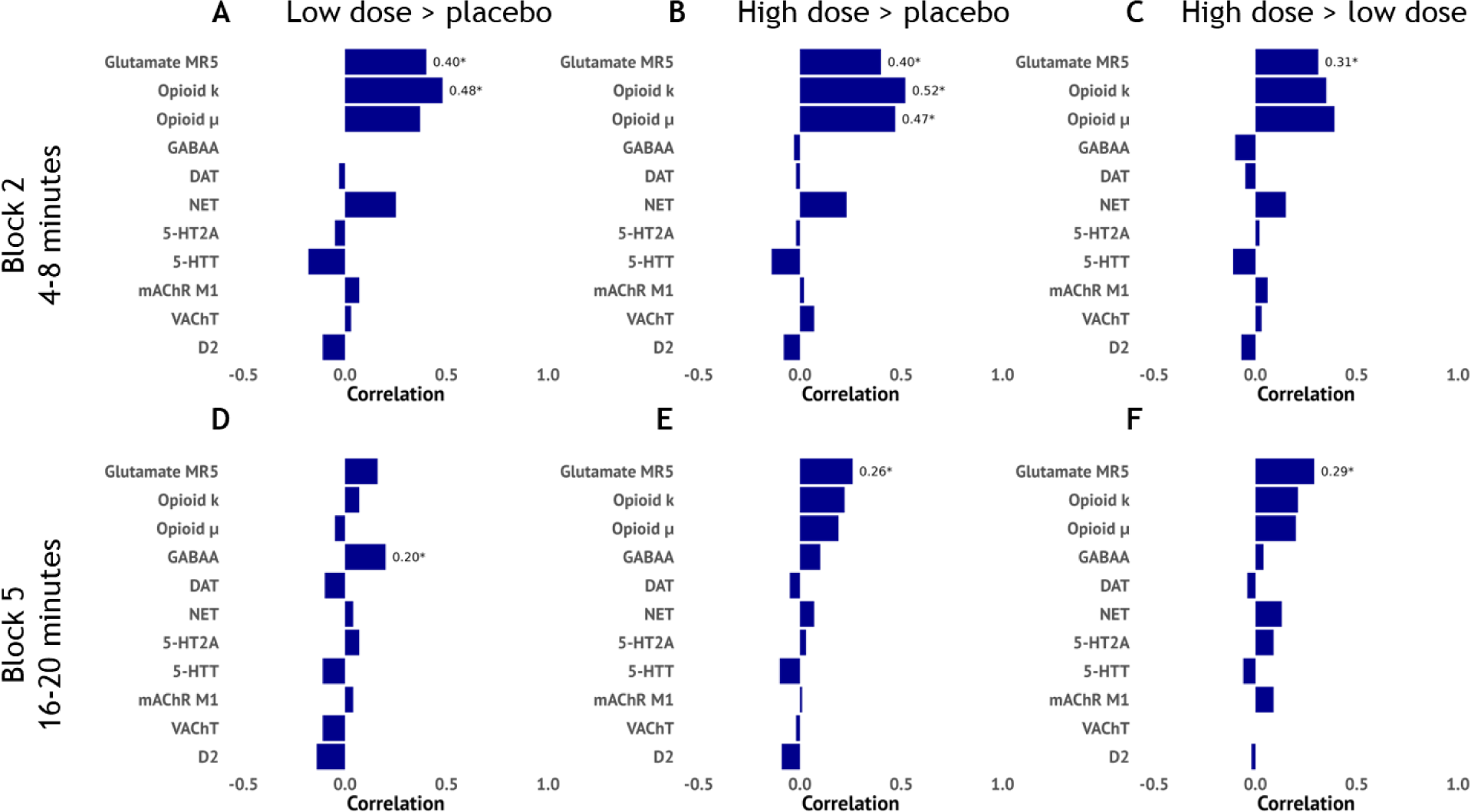
Association phMRI response and receptor distribution maps. Horizontal bar plot showing the correlation between the phMRI contrast maps and receptor distribution maps based on positron emission tomography (PET) data from healthy volunteers. These data are plotted for the low>placebo, high>placebo and high>low contrasts and for the more rapid (block 2) and slower response (block 5). Correlation coefficients were obtained using spin tests. phMRI: pharmacological magnetic resonance imaging; Glu MR5: glutamate metabotropic receptor 5; GABAA: GABAA receptor; 5-HTT: serotonin transporter; 5-HT2A: serotonin 2A receptor; VAChT: acetylcholine transporter; mAChR M1: metabotropic acetylcholine receptor 1; DAT: dopamine transporter; Dopamine D2: dopamine D2 receptor; NET: norepinephrine transporter.

## Discussion

The aim of our study was to evaluate an innovative MRI protocol capable of simultaneously assessing hemodynamic changes with phMRI and neurometabolism with phMRS to examine time- and dose-dependent responses to S-ketamine administration. This marks the first investigation employing a double-dose, placebo-controlled, within-subjects design with prolonged functional neuroimaging measurements. By integrating these two methods, we offer comprehensive and time-resolved insights into the functional dynamics induced by ketamine administration.

### phMRI response to ketamine

We observed a consistent increase in the BOLD response across various frontal, cingulate, and insular regions following both ketamine doses, aligning with findings from prior phMRI investigations involving intravenous ketamine challenges (for reviews, see^26,30^). However, our study stands out in several aspects of its design: we administered S-ketamine instead of racemic (R/S) ketamine, employed a double-dose rather than single-dose protocol, utilized a 7T-MRI scanner as opposed to lower field strengths, and extended our monitoring period to 30 minutes post-administration, a departure from the more typical 10-minute duration.

Ketamine phMRI investigations typically adopt a bolus with continuous infusion approach ^26^, although variations in administration schemes lead to differing plasma levels of ketamine. Additionally, although S-ketamine exhibits three to fourfold greater binding affinity for the NMDA receptor than R-ketamine (Ebert et al.^31^, cited in Jelen et al.^32^), direct comparisons of phMRI responses between S-ketamine, R-ketamine and racemic ketamine are lacking. Previous studies employing either racemic ketamine^13,14,17,22,33–35^ or S-ketamine^36^ have demonstrated significant activation of the ACC, insula, and thalamus across a wide range of sub-anesthetic doses. The S-ketamine dose used by Hoflich et al.^36^, which closely resembles our low-dose infusion regimen, elicited significantly weaker and less widespread activation patterns compared to our results, despite using a signal model, which should provide adequate power. These differences support the advantage of employing higher field strength acquisitions to enhance the SNR, but may additionally be influenced by differences in analytical approaches.

Although we observed significant differences in the extent of dissociation between the low and high doses, these distinctions were not mirrored in significant differences in the phMRI response. In the only other study that used multiple doses, the authors also did not report voxel-wise differences between their low and high subanesthetic doses of ketamine between subjects. These findings suggest that even relatively low doses can evoke substantial and widespread functional brain responses, with these hemodynamic effects already exhibiting ceiling effects at subanesthetic doses. In task-fMRI studies, low to moderate levels of sensory stimuli generally cause a linear increase in the BOLD response. However, the response can saturate, leading to a plateau^37^. This has been observed across sensory modalities, including visual, auditory, and somatosensory systems. Similarly, such a ceiling effect can be observed with ketamine administration. This is important to consider when interpreting phMRI results from strong pharmacological stimuli and emphasizes the importance of using complementary techniques that more directly reflect neuronal activation, like phMRS.

Most studies examining the acute effects of ketamine have primarily measured the BOLD response over relatively short periods, providing limited insight into the prolonged response. Notably, Hoflich et al.^36^ conducted one of the few studies extending this observation window to approximately 45 minutes post-bolus infusion. However, their approach solely incorporated the pharmacokinetic curve, potentially overlooking extended effects on the BOLD response. In contrast, our pseudo-block design facilitated evaluation of the phMRI response across multiple discrete bins. Intriguingly, following the peak response, we noted a transient signal decline to baseline, succeeded by a gradual BOLD signal increase in the high dose. While the origin of these temporal patterns remains speculative, evidence from electroencephalography (EEG) and magnetic encephalography (MEG) studies suggests the existence of multiple time courses of oscillatory activity with distinctive spatial profiles^38^.

When assessing the correlation between the phMRI response maps and receptor distributions, we observed that both the rapid and sustained functional responses to ketamine are associated mainly with the glutamate MR5 receptor map, consistent with ketamine’s direct effect on glutamate release^39^. Furthermore, only the rapid response to S-ketamine was associated with κ and μ receptor maps in a dose-dependent manner. Interestingly, the interaction of ketamine with both μ and κ opioid receptors appears necessary for its antidepressant properties, as illustrated by the attenuation of ketamine’s antidepressant effect following treatment with opioid receptor antagonists^40,41^. Ketamine is currently also used as analgesic and anti-hyperalgesic in subanesthetic doses as used in this study, which is thought to rely on both NMDA and μ opioid receptor binding^42^. Indeed, the observed association with the μ opioid receptor at the high dose aligns with the doses used for pain relief, supporting the known pharmacotherapeutic properties of ketamine.

### Glutamatergic origin of phMRI changes: a role for phMRS

Ketamine’s primary mechanism of action is through N-methyl-D-aspartate (NMDA) receptor antagonism^26^. It binds to the phencyclidine binding site with a slow off rate, resulting in prolonged tonic blockade of NMDA receptor currents^43^. Ketamine is believed to selectively antagonize GABA-ergic inhibitory interneurons^44^, thereby disrupting their tonic inhibitory input to cortical pyramidal neurons. This leads to a surge in glutamate release from pyramidal neurons, subsequently affecting various post-synaptic receptors, including glutamatergic AMPA receptors, as well as cholinergic and monoaminergic receptors^45,46^.

Pre-treatment studies with e.g. metabotropic glutamate 2/3 receptor agonists and lamotrigine, a voltage-gated sodium channel blocker, have demonstrated attenuation of the BOLD signal by these drugs, supporting the involvement of a ketamine-induced surge of glutamate in the cortical increases in BOLD signal seen following ketamine infusion^33,35^. Yet, to directly investigate neurometabolites involved in neuronal activity and neurotransmitter cycling, MRS is currently the only non-invasive neuroimaging technique available. We recently reviewed the studies that investigated effects of acute ketamine on neurometabolite levels^20^, and found quite mixed results. Although a few studies noted increased Glu, Gln or Glx in ACC regions^22,34,47^, the majority of studies at either 3T or 7T did not report ketamine-induced changes therein. These inconsistencies may be related to differences in study design, the variation in voxel placement and other acquisition parameters, as well as analysis choices.

All prior ketamine MRS studies have analyzed neurometabolite levels as static signals (mostly at one time-point post-administration), providing a snapshot of neurometabolite levels following ketamine administration. However, this approach foregoes the dynamic nature of neurometabolite concentrations, as has repeatedly been shown in task-activation fMRS studies^6,8,48^. As the first phMRS study taking a dynamic analysis approach, we could leverage the increased sensitivity to demonstrate time-dependent effects of ketamine administration on multiple neurometabolites. We found that particularly the high dose S-ketamine increased glutamate between approximately 8 to 32 minutes following bolus administration, which reflects a different time course compared to the phMRI response, which seems to peak earlier. In addition to glutamate, we also detect elevated lactate levels, which have been linked to non-oxidative metabolism and have been previously observed in task-related fMRS studies^10^. This finding is noteworthy, as it aligns with PET studies demonstrating increased CBF and cerebral metabolic rate of glucose in the ACC, despite minimal changes in cerebral metabolic rate of oxygen^49^. Together, this supports the hypothesis that non-oxidative glucose metabolism may be occurring during the acute hyperglutamatergic state induced by ketamine. Finally, we observed alterations in neurometabolites like tNAA and tCr, which are less commonly associated with neural activity changes. These alterations might be explained by changes in diffusion or neuronal microstructure^50,51^ or by incomplete correction for BOLD-induced effects on metabolite linewidths, as they represent the largest peaks in the spectrum^6^. However, the exact nature of these changes and the role of dynamic fitting choices herein remains to be investigated.

### Application and methodological considerations

Incorporating phMRS acquisitions into phMRI protocols combines the strengths of both techniques: the spatial resolution and SNR of fMRI acquisitions and the neural specificity of MRS. In addition to pre-treatment studies and receptor-informed fMRI, phMRS offers a unique opportunity to elucidate the neural mechanisms underlying the phMRI response. For instance, in the context of ketamine, our protocol can potentially uncover functional brain responses associated with psychosis sensitivity, as indicated by our associations with dissociation. Additionally, this approach holds promise for investigating mechanisms related to treatment response in conditions such as treatment-resistant depression. Furthermore, this protocol could be valuable to assess drug responses in populations with compromised neurovascular coupling, such as individuals with vascular pathology, where altered hemodynamic responses occur due to factors like vascular damage or arterial stiffness. In such cases, phMRS can assess the extent to which the neural response to medication remains intact. Moreover, the advantage of the simultaneous assessment of phMRI and phMRS is significant, as this reduces the risk of different outcomes due to habituation or sensitization that might occur if they were acquired on separate days. Additionally, our extended acquisition time enables the observation of effects over a longer time span compared to previous studies, potentially reflecting the activation of different receptors and downstream mechanisms.

We need to acknowledge several limitations of this study. Firstly, due to the significantly lower concentrations of neurometabolites compared to water, phMRS suffers from limited SNR in comparison to phMRI. However, our transition to 7T MRI has provided a substantial increase in SNR, enabling us to reliably estimate concentrations of several small metabolites. Additionally, through the implementation of a dynamic shimming approach^52^, we achieved good quality spectra and phMRI data with our interleaved protocol. Despite these improvements, we encountered challenges in dynamically fitting GABA and Gln resonances. Addressing this in future studies would be valuable, as these metabolites are closely linked to glutamate and could offer insights into the excitation-inhibition balance, potentially elucidating the phMRI results further. Furthermore, the temporal resolution of our interleaved acquisitions is predominantly dictated by the adiabatic pulses in the sLASER sequence, suggesting the need for future studies to explore methods for accelerating these acquisitions.

## Conclusion

Our study demonstrates a robust phMRI response in the dorsal-frontal, cingulate, and insular cortices for both low and high subanesthetic doses of S-ketamine compared to placebo. The phMRI response patterns closely align with the distribution of glutamate and opioid receptors. Additionally, these hemodynamic changes coincide with increased glutamate and lactate levels in the ACC for the high dose. Together, simultaneous phMRI and phMRS measurements offer a more comprehensive understanding of the brain’s functional response to S-ketamine, as demonstrated by the improved accuracy in predicting S-ketamine doses and placebo conditions when combining the two techniques. Our novel interleaved phMRI-phMRS protocol bears considerable potential for concurrently assessing hemodynamic and neurometabolic effects of ketamine and other pharmacological agents. This innovative approach provides a pathway to disentangle the contributions of various neurometabolic processes to medication-induced functional brain changes. Moving forward, it holds great promise for enhancing our understanding of drug-brain interactions and facilitating a more direct investigation into the neurometabolic mechanisms underlying therapeutic response.

## Methods

The study was registered in the Dutch Trial Register ([NL8994](https://onderzoekmetmensen.nl/nl)). A pre-registration for analyses was done at Open-Science Framework (https://osf.io/6hzte). All procedures were approved by the medical ethics committee (NL74447.018.20) of the Amsterdam University Medical Center in Amsterdam prior to the start of the study and were conducted in accordance with the Declaration of Helsinki. All subjects gave written informed consent.

### 2.1 Participants

We recruited healthy volunteers with the following inclusion criteria: 18-55 years of age BMI between 18.5 and 25; hormonal contraception for females (no hormone-free days during the study). Participants with a (history of) psychiatric treatment, having first-degree relatives with (a history of) schizophrenia and major depression disorder, with (a history of) neurological disorders or concussion with loss of consciousness were excluded. Additionally, we excluded participants with contraindications for S-ketamine administration or 7T MRI, and with (a history of) drug or alcohol dependence. Participants were recruited via flyers and subject recruitment websites.

### 2.2 Study design

In total, participants visited the laboratory site four times. During the first visit, participants were screened for (history of) psychiatric disorders (Mini International Neuropsychiatric Interview, M.I.N.I)), alcohol or drug dependence (Alcohol, Smoking, and Substance Involvement Screening Test, ASSIST-Lite), underwent a physical examination during which height, weight, heart rate and blood pressure were measured.

During three experimental sessions, participants received placebo, a low, or a high subanesthetic dose of S-ketamine intravenously in a single-blind, double-dose, counterbalanced, placebo-controlled randomized crossover design. No feedback about the order of administration was provided until completion of the study. During each experimental session, subjects were screened using a urine test for recreational drug use and for pregnancy (women). An intravenous line was placed by a resident anesthesiologist for S-ketamine administration and blood sampling. After ∼20 minutes of baseline MRI-scans, the administration of S-ketamine or placebo commenced using an infusion pump. The administration consisted of a bolus (1 min), followed by a continuous administration (∼40 min) to keep plasma levels stable throughout the remainder of the scans (placebo: NaCl 0.9%; low dose: 0.11 mg/kg + 0.1 mg/kg/h; high dose: 0.22 mg/kg + 0.2 mg/kg/h). The dose was determined based on prior studies^26^. To monitor subjective effects of S-ketamine, a visual analog scale (VAS)^53^ and the Clinician-Administered Dissociative States Scale (CADSS)^54^ were administered four times: at the start and end of each experimental session, and two times in the MRI-scanner: before the start of the administration, and immediately after termination of the administration (participants were instructed to report on experienced dissociation during the administration period). Following the MRI scan, a blood sample was collected by an anaesthesiologist to determine S-ketamine and nor-S-ketamine blood plasma concentrations.

Participants were instructed to refrain from using psychotropic medication and recreational drug use over a period of 1 week prior to each session and from using alcohol 24 hours before each session. On the day of the session, participants were instructed to refrain from caffeine. All sessions for each participant were scheduled at the same time of day (max 1-hour difference between sessions), to minimize effects of circadian rhythm on the outcomes of the study. Study days were separated by at least 7 days.

### 2.3 Acquisition

Data was acquired on a 7T MR system (Philips, Best, the Netherlands) with a dual-channel transmit and 32-channel receive head coil (Nova Medical Inc., Wilmington, MA, USA). A T1-weighted (T1w) structural scan was obtained (TR/TE/FA = 5ms/2ms/7°; FOV(AP,FH,RL)=246x246×180 mm3; voxel size=0.85×0.85×1 mm3), followed by an interleaved fMRI/fMRS protocol^8^ (dynamic scan time 5.1 seconds; dynamic linear shim and shared optimized set of second order shims^52^; total scan duration 38 min). The fMRI sequence was a 3D-EPI: TR/TE/FA=31/20ms/10∘; voxel size=1.875×1.875×2.09 mm3; FOV=240x240x136.5 mm3. In addition, a topup scan was obtained for distortion correction with the same parameters, but opposing phase encoding direction (4 volumes). For MRS a semi-LASER sequence was used: TR/TE=3500/36ms; bandwidth=3kHz; number of points=1024; ACC volume of interest (VOI) size=30x20x15 mm3; VAPOR water-suppression. The VOI was placed in the dorsal ACC (Fig. 1B). Participants were instructed to focus on a fixation cross and let their minds wander.

### 2.4 Processing and analysis

#### 2.4.1 phMRI

PhMRI data were preprocessed using in-house scripts and fMRIprep v20.1.1^55^. which is based on Nipype 1.4.2^56^. In short, T1w scans were normalized to Montreal Neurological Institution (MNI) space. Motion correction (FLIRT), distortion correction (fieldmap), and T1w co-registration were performed as part of the functional data preprocessing. Then, the first 12 volumes were discarded, and WM, CSF, and 6 motion parameters (rotation and translation in x, y, and z direction) and their first derivatives were regressed from the signal, followed by spatial smoothing (4mm FWHM). Volumes with FD>1mm were identified. First-level GLM analyses contrasted the administration blocks to baseline, yielding 8 first-level contrasts per subject per session, and were carried out in FSL-FEAT^57,58^ (Supplementary Fig. S4). In short, the first-level design served to divide the data in 9 blocks of 48 volumes (∼4 minutes), i.e. a pseudo-block design (similar to Deakin et al.^14^; McMillan et al.^13^). The first 48 volumes modeled the baseline, where no medication was infused. The second 48 volumes modeled the bolus administration and the consecutive 7x48 volumes modeled the continuous administration in seven blocks. Higher-level analyses were set up to determine the voxel-wise time- and dose-dependent responses to placebo and S-ketamine.

#### 2.4.2 phMRS

phMRS data processing and dynamic fitting was done in FSL-MRS^23^. Through a reduction in the number of variables, and better estimation of highly correlated model terms, dynamic fitting approaches have been shown to enhance sensitivity (precision) in detecting stimulus-induced shifts in neurometabolite levels compared to block averaging^24,59^. Processing included coil combination, phase-and frequency alignment, and eddy-current correction. The first 14 and last four transients were removed to match the number of transients to the fMRI volumes.

For quality control, the remaining transients for each subject were combined and averaged, and the resulting average spectrum underwent fitting using FSL-MRS (version 2.1.13) to assess spectral quality. This fitting process involved aligning basis spectra to the complex-valued spectrum in the frequency domain. A basis set was simulated based on our MRS sequence in FSL-MRS. The basis set was simulated using full pulse descriptions and 60 spatially resolved points in each dimension^60,61^. The basis set comprised 19 metabolites and a measured macromolecular baseline, available at https://github.com/mrshub/mm-consensus-data-collection^62^. A Voigt lineshape model, incorporating one Gaussian and one Lorentzian line broadening parameter, was applied to shift and broaden the basis spectra. Metabolite basis spectra shared the same lineshape model and parameters, except for macromolecules, which had distinct parameters. Specifically, the lineshape model employed four linewidth parameters: one Lorentzian and one Gaussian parameter for all metabolites, and another set for the macromolecular (MM) basis spectrum. The Lorentzian parameter was enforced to be positive, as was the constant term of the Gaussian broadening. Additionally, a complex second-order polynomial baseline was concurrently fitted. Model fitting utilized the truncated Newton algorithm from scipy^63^. Spectra were excluded for poor spectral quality if Cramer-Rao Lower Bounds (CRLB) exceeded 5% for Glu, and the full-width-half-maximum (FWHM) of the linewidth of total Creatine (tCr) exceeded 19Hz. Spectral quality metrics are shown in Supplementary Table S20.

For the time series analysis, all data underwent dynamic fitting within FSL-MRS, enabling simultaneous fitting of a dynamic signal model to all transients. This approach, also known as spectral-temporal fitting or 2D fitting^59^, utilized a design matrix to model the temporal response of each metabolite in the context of a GLM analysis. This matrix, combined with linear combination modeling of the spectral response at each timepoint, was fitted to the entire dataset concurrently using a least-squares approach. Spectral fitting parameters remained consistent with those described in the paragraph above. In this dynamic spectral fitting model, various parameters such as phase, shift, baseline, linewidth, and concentration could be time-dependent variables. While the phase, shift, and baseline parameters were fixed across all timepoints, the concentration and linewidth were constrained by the temporal model (design matrix). Notably, the linewidth was varied, considering previous findings suggesting stimulus-induced changes in the BOLD effect could lead to line narrowing in MR spectra due to alterations in T2*^6,25^. Note that we applied the design matrix to the Gaussian line-broadening parameter, whereas the Lorentzian parameter remained fixed across time.

Design matrices, created using nilearn^64^, incorporated separate regressors for each of the blocks, similar to the phMRI matrix, as well as a baseline regressor. The design matrix fitted to the concentration and linewidth were the same, except that a linear drift parameter was added to the linewidth design matrix (to reflect potential drift in linewidth as a result of BOLD drift) (Supplementary Fig. S4).

Acquisition and analysis parameters for fMRS were reported following the MRSinMRS checklist^65^ (Supplementary Table S21).

### 2.5 Statistics

Higher-level analyses investigated changes from baseline for each administration block (time-dependent effects) and condition differences in these changes from baseline per administration block (dose-dependent effects) for both the phMRS and phMRI data.

#### 2.5.1 phMRI

For the phMRI data, both time- and dose-dependent effects were determined in FSL Permutation-based Analysis of Linear Models^66^, which allows for voxel-wise correction for multiple testing. For both time- and dose-dependent analyses, we used 5000 permutations and threshold-free cluster enhancement^67^. Significance was inferred when family-wise error corrected p-values (p_FWE_) <0.05; p_FWE_ values were corrected for multiple comparisons within each contrast image, as well as for multiple testing across modalities (administration-baseline contrasts). Time-dependent effects per condition were determined using one-sample t-tests per administration-baseline contrast; dose-dependent effects were determined using paired t-tests. Percentage overlap between significant clusters and Daemon (standard Tailararch) atlas regions for each of these comparisons were determined using *whereami* from AFNI^68,69^.

#### 2.5.2 phMRS

Likewise, for the phMRS data, time-dependent effects per condition were determined using one-sample t-tests per administration-baseline contrast using FSL-MRS. Dose-dependent effects between conditions for each administration-baseline contrast were conducted using paired t-tests in FSL-MRS, using FSL’s *flameo*^58^. Multiple comparison correction was performed for both time- and dose-dependent effects using False Discovery Rate (FDR) (stats package (R Core Team, 2013)), with significance inferred when p_FDR_ <0.05. Metabolites were excluded from further fMRS analyses if the CRLB exceeded 20% in >50% of the participants based on the average spectrum. In addition, time- and dose-dependent effects of S-ketamine on Gaussian linewidth (sigma) were investigated, as decreases in sigma have been shown to be associated with stimulus-induced increases in BOLD^6^.

#### 2.5.3 Association phMRI and receptor distribution maps

Receptor distribution maps were obtained from neuromaps^27^, an open source toolbox accessing, transforming and analyzing structural and functional brain annotations. All receptor maps of interest were available via neuromaps and are provided in Table 1 below. All maps were resampled to 2mm isotropic resolution and parcellated into 250 atlas regions, based on the Cammoun atlas available in neuromaps^70^. To quantify the association between phMRI response maps and various receptor distributions, we performed a spin-test^27,28^, a statistical method employed to compute null models by randomly rotating data, allowing for the assessment of statistical significance between two spatial maps^29^. Spatial null distributions were generated for each of the investigated placebo-drug-contrast based on whole-brain t-stat maps by applying the approach by Burt et al.^71^ (default parameters as provided by the brainSMASH software; 1000 permutations) and were subsequently correlated to each of the parcellated receptor maps. Due to the exploratory nature of these analyses, we report the uncorrected p-values per association.

**Table 1.**
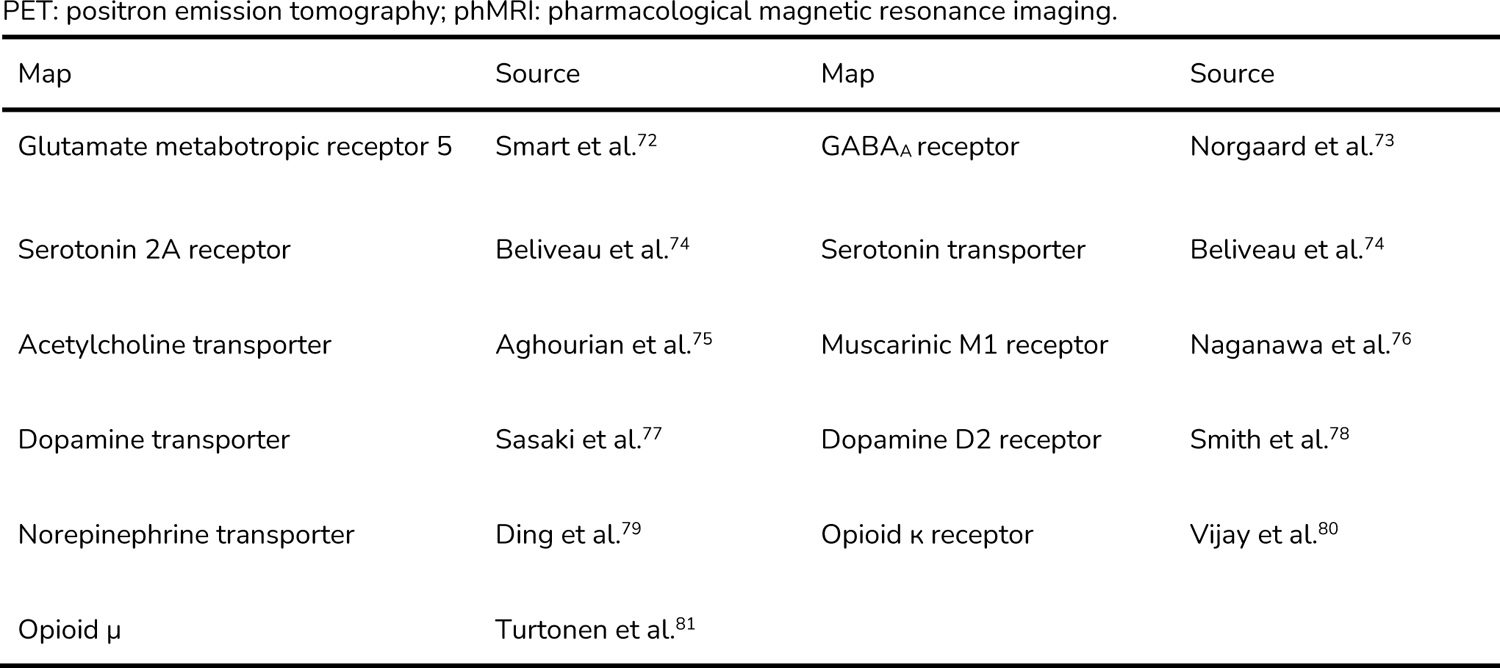
Overview of used PET maps to investigate the association between the phMRI response and receptor distributions.

PET: positron emission tomography; pHMRI: pharmacological magnetic resonance imaging.
| Map | Source | Map | Source |
| --- | --- | --- | --- |
| Glutamate metabotropic receptor 5 | Smart et al.<sup>72</sup> | GABA<sub>A</sub> receptor | Norgaard et al.<sup>73</sup> |
| Serotonin 2A receptor | Beliveau et al.<sup>74</sup> | Serotonin transporter | Beliveau et al.<sup>74</sup> |
| Acetylcholine transporter | Aghourian et al.<sup>75</sup> | Muscarinic M1 receptor | Naganawa et al.<sup>76</sup> |
| Dopamine transporter | Sasaki et al.<sup>77</sup> | Dopamine D2 receptor | Smith et al.<sup>78</sup> |
| Norepinephrine transporter | Ding et al.<sup>79</sup> | Opioid $\kappa$ receptor | Vijay et al.<sup>80</sup> |
| Opioid $\mu$ | Turtonen et al.<sup>81</sup> | | |

#### 2.5.4 Statistical analyses

Subject-level BOLD COPE values from phMRI analyses were extracted from the ACC phMRS voxel using *fslstats*. COPE values were analyzed using linear mixed-effects models in RStudio (package nmle^82,83^ to determine both time- and dose-dependent effects (random intercept for participant to account for repeated measures, fitted using the maximum log-likelihood estimation; ML). The influence of covariates (i.e. order of S-ketamine administration, mean FD, %change in heart rate from baseline) was assessed to determine the stability of these findings. Model selection for all models was conducted as follows: the most complicated model in terms of main predictors and interaction effects was tested at first, and subsequently predictors not significantly contributing to the prediction were removed from the model. The final model included only the remaining significant interaction effects, significant main effects, and main effects of predictors for the significant interaction effects. Each final model with the outcome tables is reported in the Supplementary Results. Differences in blood plasma levels of S-ketamine and nor-S-ketamine between the low and high S-ketamine dose conditions were analyzed using Wilcoxon signed rank tests. Subjective dissociation was analyzed using Friedman’s test on post-administration total CADSS scores, followed by Wilcoxon signed rank tests to determine differences between conditions. Significance was inferred when p_FDR_<0.05.

Bayesian multilevel modeling (*package brms*^84–86^; a high-level interface of *stan*^87^), was used to predict condition (placebo, low S-ketamine dose, high S-ketamine dose) based on individual-level COPE values. These regression models have the ability to account for the repeated measured structure of the data. All models were specified with four Markov chains, each with 2000 iterations, including 1000 warmup iterations. Three different model types were specified: I. phMRI COPE values only, II. phMRS COPE values only, and III. COPE values from both phMRI and phMRS. Per model type, different models were specified (random intercept for participant, random intercept and slope per participant) and were compared using the Leave-One-Out Cross-Validation (LOO) criterion and the Widely Applicable Information Criterion (WAIC) to evaluate their predictive performance (*package loo*^88^). Per model type (I, II, III), the best-fitting model was selected and model fit was once again compared using LOO-IC and WAIC. All Bayesian models were fitted on subjects with complete datasets for all predictors (N=23).

All statistical analyses were conducted in RStudio v2022.02.1 (RStudio Team (2020); R version 4.3.1).

