## Supplementary Materials for "Concurrent assessment of neurometabolism and brain hemodynamics to comprehensively characterize the functional brain response to psychotropic drugs: an S-ketamine study" (from another version of this preprint: version 1, 2024-07-24)

Supplementary Methods ... 3
Preprocessing of functional MRI data using fMRIprep ... 3
Anatomical data preprocessing ... 3
Functional data preprocessing ... 3
References ... 4
Supplementary Results ... 7
Time-dependent changes in BOLD ... 7
Time-dependent changes in neurometabolite levels ... 7
Supplementary Figures ... 8
Fig. S1. Time-dependent effects of S-ketamine on whole-brain fMRI BOLD ... 8
Fig. S2. Analysis of phMRI data using a signal model analysis. ... 8
Table S1. Cluster information for time-dependent effects of S-ketamine on the phMRI signal for the placebo condition. ... 10
Table S2. Cluster information for time-dependent effects of S-ketamine on the phMRI signal for the low S-ketamine dose condition ... 12
Table S3. Cluster information for time-dependent effects of S-ketamine on the phMRI signal for the high S-ketamine dose condition ... 13
Table S4. Cluster information for dose-dependent effects of S-ketamine on the phMRI signal... 15
Table S5. phMRI response: Linear mixed effect model for individual-level COPE values in the ACC MRS voxel. ... 17
Table S6. phMRI response and heart rate: Linear mixed effect model for individual-level COPE values in the ACC MRS voxel. ... 18
Table S7. phMRI response and framewise displacement (FD): Linear mixed effect model for individual-level COPE values in the ACC MRS voxel ... 20
Table S8. phMRI response and medication order: Linear mixed effect model for individual-level COPE values in the ACC MRS voxel. A. ... 21
Table S9. Heart rate: linear mixed effect model for %change in heart rate from baseline. ... 22

### Supplementary Methods

#### Preprocessing of functional MRI data using fMRIprep

Results included in this manuscript come from preprocessing performed using fMRIPrep 20.1.1 (Esteban, Markiewicz, et al. (2018); Esteban, Blair, et al. (2018); RRID:SCR_016216), which is based on Nipype 1.5.0 (Gorgolewski et al. (2011); Gorgolewski et al. (2018); RRID:SCR_002502).

**Anatomical data preprocessing**

A total of 3 T1-weighted (T1w) images were found within the input BIDS dataset. All of them were corrected for intensity non-uniformity (INU) with N4BiasFieldCorrection (Tustison et al. 2010), distributed with ANTs 2.2.0 (Avants et al. 2008, RRID:SCR_004757). The T1w-reference was then skull-stripped with a Nipype implementation of the antsBrainExtraction.sh workflow (from ANTs), using OASIS30ANTs as target template. Brain tissue segmentation of cerebrospinal fluid (CSF), white-matter (WM) and gray-matter (GM) was performed on the brain-extracted T1w using fast (FSL 5.0.9, RRID:SCR_002823, Zhang, Brady, and Smith 2001). A T1w-reference map was computed after registration of 3 T1w images (after INU-correction) using mri_robust_template (FreeSurfer 6.0.1, Reuter, Rosas, and Fischl 2010). Volume-based spatial normalization to two standard spaces (MNI152NLin2009cAsym, MNI152NLin6Asym) was performed through nonlinear registration with antsRegistration (ANTs 2.2.0), using brain-extracted versions of both T1w reference and the T1w template. The following templates were selected for spatial normalization: ICBM 152 Nonlinear Asymmetrical template version 2009c [Fonov et al. (2009), RRID:SCR_008796; TemplateFlow ID: MNI152NLin2009cAsym], FSL's MNI ICBM 152 non-linear 6th Generation Asymmetric Average Brain Stereotaxic Registration Model [Evans et al. (2012), RRID:SCR_002823; TemplateFlow ID: MNI152NLin6Asym],

**Functional data preprocessing**

For each of the 3 BOLD runs found per subject (across all tasks and sessions), the following preprocessing was performed. First, a reference volume and its skull-stripped version were generated using a custom methodology of fMRIPrep. Head-motion parameters with respect to the BOLD reference (transformation matrices, and six corresponding rotation and translation parameters) are estimated before any spatiotemporal filtering using mcflirt (FSL 5.0.9, Jenkinson et al. 2002). A B0-nonuniformity map (or fieldmap) was estimated based on two (or more) echo-planar imaging (EPI) references with opposing phase-encoding directions, with 3dQwarp Cox and Hyde (1997) (AFNI 20160207). Based on the estimated susceptibility distortion, a corrected EPI (echo-planar imaging) reference was calculated for a more accurate co-registration with the anatomical reference. The BOLD reference was then co-registered to the T1w reference using flirt (FSL 5.0.9, Jenkinson and Smith 2001) with the boundary-based registration (Greve and Fischl 2009) cost-function. Co-registration was configured with nine degrees of freedom to account for distortions remaining in the BOLD reference. The BOLD time-series (including slice-timing correction when applied) were resampled onto their original, native space by applying a single, composite transform to correct for head-motion and susceptibility distortions. These resampled BOLD time-series will be referred to as preprocessed

BOLD in original space, or just preprocessed BOLD. The BOLD time-series were resampled into standard space, generating a preprocessed BOLD run in MNI152NLin2009cAsym space. First, a reference volume and its skull-stripped version were generated using a custom methodology of fMRIPrep. Automatic removal of motion artifacts using independent component analysis (ICA-AROMA, Pruim et al. 2015) was applied to collect the "aggressive" noise-regressors and place them in the corresponding confounds file. Several confounding time-series were calculated based on the preprocessed BOLD: framewise displacement (FD), DVARS and three region-wise global signals. FD was computed using two formulations following Power (absolute sum of relative motions, Power et al. (2014)) and Jenkinson (relative root mean square displacement between affines, Jenkinson et al. (2002)). FD and DVARS are calculated for each functional run, both using their implementations in Nipype (following the definitions by Power et al. 2014). The three global signals are extracted within the CSF, the WM, and the whole-brain masks. Additionally, a set of physiological regressors were extracted to allow for component-based noise correction (CompCor, Behzadi et al. 2007).

Many internal operations of fMRIPrep use Nilearn 0.6.2 (Abraham et al. 2014, RRID:SCR_001362), mostly within the functional processing workflow. For more details of the pipeline, see the section corresponding to workflows in fMRIPrep's documentation.

### Supplementary Results

#### Time-dependent changes in BOLD

At the voxel-wise level, both sub-anesthetic S-ketamine doses induced significant increases in the BOLD signals, compared to baseline, across multiple time points (Supplementary Figure S1). The spatial pattern of increased BOLD responses covers the medial frontal, (para)cingulate, and insular areas. The response to both S-ketamine doses was characterized by a strong early rise in the BOLD signal (0-8 minutes of administration), which subsequently normalized to baseline, but increased again after 16 minutes for the high S-ketamine dose only.

#### Time-dependent changes in neurometabolite levels

Compared to baseline, Glu significantly increased during the second ($p_{FDR}$=0.001), third ($p_{FDR}$=0.01), and fourth ($p_{FDR}$=0.01) administration blocks (4-16 minutes), respectively. No significant changes in Glu were found for the low dose compared to baseline. We observed no significant changes from baseline for aspartate for any of the conditions. For glucose, a significant decrease in glucose from baseline was observed during the high S–ketamine dose only (block 1: $p_{FDR}$=0.01; block 2: $p_{FDR}$=0.01; block 4: $p_{FDR}$=0.048). Compared to baseline, lactate increased 4-12 min post-administration in the low dose (block 2: $p_{FDR}$=0.005; block 3: $p_{FDR}$=0.048) and in the first 28 minutes post-administration for the high S-ketamine dose (block 1: $p_{FDR}$=0.03; block 2: $p_{FDR}$<0.0001; block 3: $p_{FDR}$=0.002; block 4: $p_{FDR}$=0.003; block 5: $p_{FDR}$=0.004; block 6: $p_{FDR}$=0.03; block 7: $p_{FDR}$=0.02).

Gaussian lineshape parameter sigma decreased during the first administration block in the low condition ($p_{FDR}$=0.02) and during the first two post-administration blocks in the high condition (block 1: $p_{FDR}$<0.0001; block 2: $p_{FDR}$=0.0003), in line with the phMRI findings in the ACC voxel.

Time- and dose-dependent changes in other neurometabolites are more difficult to interpret, as they are not as explicitly involved in the neurometabolic cycle or have not been extensively assessed in fMRS studies previously. For example, we observe changes in PE, Ins, tNAA, tCr, and tCh. PE increased significantly in the low dose compared to placebo, whereas Ins increased significantly in the high dose compared to placebo. Additionally, significant differences in tNAA, tCr and tCh between the conditions were observed (Supplementary Tables S13-14).

#### Supplementary Figures

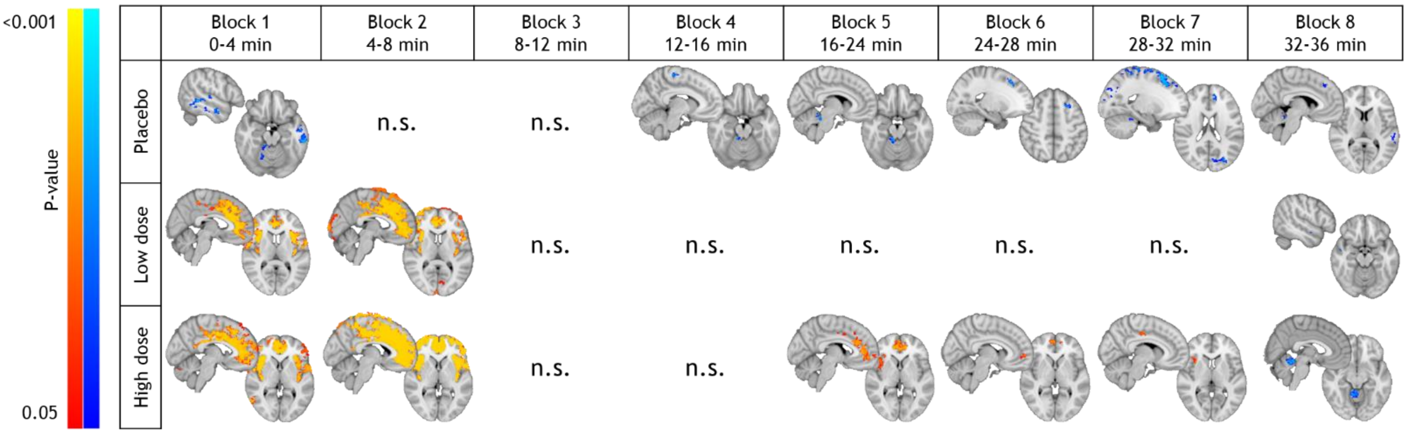

**Fig. S1. Time-dependent effects of S-ketamine on whole-brain fMRI BOLD**. Cluster information from the voxel-based analysis showing changes from baseline for each condition (placebo, low dose of S-ketamine, and high dose of S-ketamine) per block.

BOLD: blood-oxygen level dependent; phMRI: pharmacological magnetic resonance imaging.

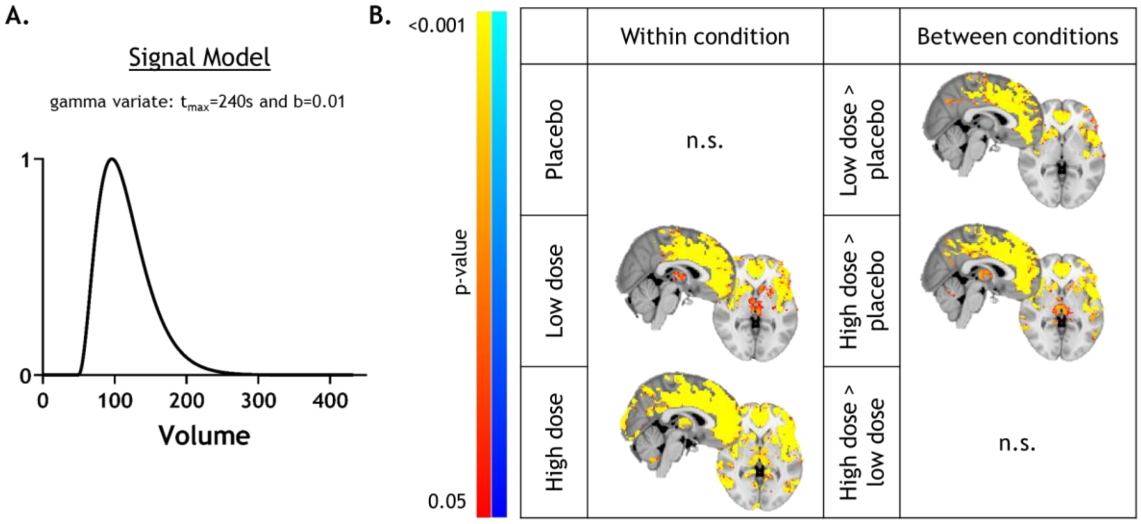

**Fig. S2. Analysis of phMRI data using a signal model analysis. A.** The used signal model regressor. We modeled this regressor as a gamma variate using $t_{max}$=240s and b=0.01. **B**. Main effects of the signal model regressor per condition are depicted on the left (within condition). Dose-dependent effects are visualized on the right (between conditions).

phMRI: pharmacological magnetic resonance imaging.

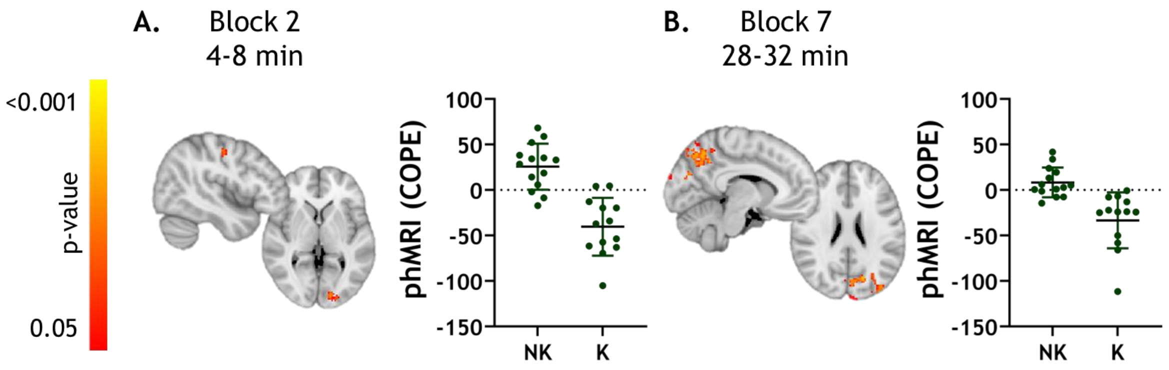

**Fig. S3. Effect of prior ketamine use on placebo response.** Group differences between subjects who had experienced ketamine prior to the experiments and those who had not in the placebo session. **A.** Significant differences between the two groups during infusion block 2. **B.** Significant differences between the two groups during infusion block 7.

COPE = contrast of parameter estimates; K= prior ketamine experience; NK= no prior ketamine experience; phMRI: pharmacological magnetic resonance imaging.

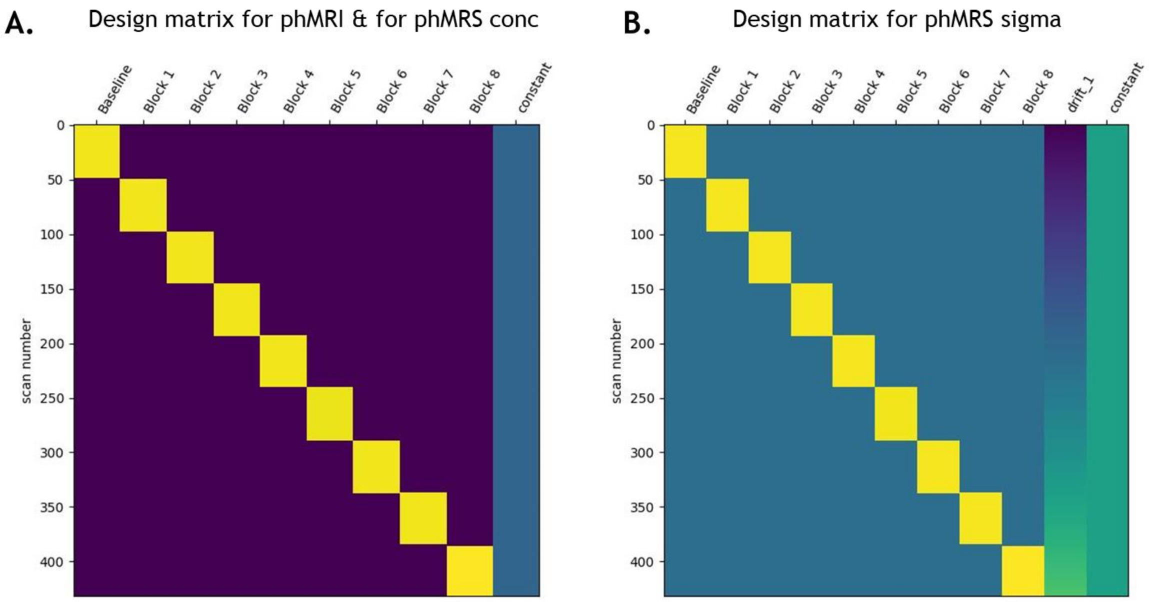

**Fig. S4. Design matrix for phMRI and phMRS. A.** The design matrix used for the first-level phMRI analysis and for the *concentration* parameter in the first-level phMRS analysis. **B.** The design matrix used for the *sigma* parameter in the first-level phMRS analysis.

conc: concentration; phMRI: pharmacological magnetic resonance imaging; phMRS: pharmacological magnetic resonance spectroscopy.

**Table S1. Cluster information for time-dependent effects of S-ketamine on the phMRI signal for the placebo condition.** Only clusters showing more than 5% overlap with an atlas region are shown. The Daemon (standard Talaraich) atlas was used identification and clusters were extracted using whereami in AFNI.

phMRI: pharmacological magnetic resonance imaging.

| Block | Cluster | # voxels | p-value | Overlap (%) | Region |
|---|---|---|---|---|---|
| 1 | 1 | 7 | .043 | 100.0 | Left Posterior Cingulate |
| | 2 | 21 | .041 | 54.8 | Left Superior Temporal Gyrus |
| | | | | 39.4 | Left Transverse Temporal Gyrus |
| | 3 | 23 | .035 | 82.0 | Right Declive |
| | | | | 14.5 | Right Culmen |
| | 4 | 160 | .017 | 63.3 | Left Inferior Temporal Gyrus |
| | | | | 19.7 | Left Middle Temporal Gyrus |
| | | | | 3.1 | Left Fusiform Gyrus |
| | 5 | 177 | .026 | 78.8 | Right Culmen |
| | | | | 9.4 | Right Cerebellar Lingual |
| | | | | 1.8 | Right Parahippocampal Gyrus |
| | 6 | 502 | .016 | 48.7 | Left Middle Temporal Gyrus |
| | | | | 18.0 | Left Superior Temporal Gyrus |
| | | | | 12.1 | Left Inferior Temporal Gyrus |
| | | | | 7.3 | Left Fusiform Gyrus |
| 4 | 1 | 16 | .034 | 100.0 | Right Culmen |
| | 2 | 45 | .033 | 55.9 | Left Medial Frontal Gyrus |
| | | | | 9.1 | Left Precentral Gyrus |
| | | | | 8.2 | Left Paracentral Lobule |
| 5 | 1 | 78 | .023 | 63.8 | Right Culmen |
| | | | | 9.9 | Right Cerebellar Lingual |
| 6 | 1 | 10 | .045 | 85.1 | Left Middle Frontal Gyrus |
| | | | | 14.9 | Left Superior Frontal Gyrus |
| | 2 | 15 | .042 | 100.0 | Left Superior Frontal Gyrus |
| | 3 | 51 | .028 | 62.3 | Left Superior Frontal Gyrus |
| | | | | 37.7 | Left Medial Frontal Gyrus |
| | 4 | 140 | .024 | 52.5 | Left Superior Frontal Gyrus |
| | | | | 44.8 | Left Middle Frontal Gyrus |
| 7 | 1 | 5 | .045 | 100.0 | Left Parahippocampal Gyrus |
| | 2 | 5 | .039 | 44.7 | Left Fusiform Gyrus |
| | | | | 31.9 | Left Culmen |
| | | | | 23.4 | Left Declive |
| | 3 | 9 | .047 | 100.0 | Left Precuneus |
| | 4 | 9 | .046 | 100.0 | Left Precuneus |
| | 5 | 9 | .045 | 53.5 | Left Postcentral Gyrus |
| | 6 | 10 | .039 | 46.5 | Left Inferior Parietal Lobule |
| | | | | 100.0 | Left Precuneus |
| | 7 | 11 | .043 | 40.8 | Left Middle Temporal Gyrus |
| | 8 | 12 | .038 | 78.3 | Left Culmen |
| | | | | 21.7 | Left Cerebellar Lingual |

| | | | | | |
|---|---|---|---|---|---|
| | 9 | 14 | .044 | 59.0 | Left Fusiform Gyrus |
| | | | | 40.3 | Left Culmen |
| | 10 | 16 | .046 | 91.6 | Left Middle Occipital Gyrus |
| | | | | 8.4 | Left Cuneus |
| | 11 | 27 | .031 | 37.6 | Left Culmen |
| | 12 | 35 | .04 | 91.4 | Left Precuneus |
| | | | | 8.6 | Left Paracentral Lobule |
| | 13 | 37 | .029 | 54.2 | Left Postcentral Gyrus |
| | 14 | 81 | .033 | 54.3 | Right Superior Parietal Lobule |
| | | | | 33.0 | Right Precuneus |
| | 15 | 103 | .024 | 10.9 | Right Precentral Gyrus |
| | | | | 8.8 | Right Paracentral Lobule |
| | | | | 8.5 | Right Medial Frontal Gyrus |
| | 16 | 128 | .033 | 55.8 | Left Postcentral Gyrus |
| | | | | 41.6 | Left Precentral Gyrus |
| | 17 | 374 | .02 | 45.1 | Left Cuneus |
| | | | | 17.9 | Left Precuneus |
| | | | | 13.7 | Left Middle Occipital Gyrus |
| | 18 | 508 | .021 | 59.0 | Right Culmen |
| | | | | 10.4 | Right Parahippocampal Gyrus |
| | 19 | 2575 | .004 | 5.9 | Left Culmen |
| | | | | 24.8 | Left Superior Frontal Gyrus |
| | | | | 23.8 | Left Middle Frontal Gyrus |
| | | | | 11.9 | Left Medial Frontal Gyrus |
| 8 | 1 | 15 | .043 | 50.3 | Left Medial Frontal Gyrus |
| | | | | 49.7 | Left Superior Frontal Gyrus |
| | 2 | 17 | .047 | 96.9 | Left Superior Temporal Gyrus |
| | 3 | 60 | .04 | 98.5 | Left Superior Temporal Gyrus |
| | 4 | 82 | .019 | 30.6 | Right Culmen |
| | | | | 29.6 | Right Cerebellar Lingual |
| | | | | 20.3 | Left Culmen |

**Table S2. Cluster information for time-dependent effects of S-ketamine on the phMRI signal for the low S-ketamine dose condition.** Only clusters showing more than 5% overlap with an atlas region are shown. The Daemon (standard Talaraich) atlas was used identification and clusters were extracted using whereami in AFNI.

phMRI: pharmacological magnetic resonance imaging.

| Block | Cluster | # voxels | p-value | Overlap (%) | Region |
|---|---|---|---|---|---|
| 1 | 1 | 7 | .047 | 73.4 | Left Precuneus |
| | | | | 7.8 | Left Paracentral Lobule |
| | 2 | 14 | .043 | 43.6 | Left Cingulate Gyrus |
| | 3 | 107 | .025 | 87.0 | Left Middle Frontal Gyrus |
| | | | | 8.2 | Left Superior Frontal Gyrus |
| | 4 | 2578 | <.0001 | 32.8 | Left Insula |
| | | | | 16.5 | Left Inferior Frontal Gyrus |
| | | | | 9.8 | Left Inferior Parietal Lobule |
| | | | | 8.6 | Left Superior Temporal Gyrus |
| | | | | 8.4 | Left Precentral Gyrus |
| | 5 | 3463 | <.0001 | 28.6 | Right Inferior Frontal Gyrus |
| | | | | 17.8 | Right Insula |
| | | | | 15.0 | Right Middle Frontal Gyrus |
| | | | | 6.9 | Right Precentral Gyrus |
| | | | | 6.1 | Right Inferior Parietal Lobule |
| | 6 | 3896 | <.0001 | 17.9 | Right Medial Frontal Gyrus |
| | | | | 13.8 | Right Cingulate Gyrus |
| | | | | 13.5 | Right Anterior Cingulate |
| | | | | 12.5 | Left Anterior Cingulate |
| | | | | 12.3 | Left Medial Frontal Gyrus |
| | | | | 6.6 | Right Superior Frontal Gyrus |
| | | | | 6.0 | Left Cingulate Gyrus |
| 2 | 1 | 5 | .047 | 97.9 | Right Cuneus |
| | 2 | 10 | .046 | 100.0 | Left Precentral Gyrus |
| | 3 | 22 | .041 | 85.6 | Left Middle Temporal Gyrus |
| | 4 | 57 | .036 | 50.8 | Right Cuneus |
| | | | | 28.7 | Right Superior Occipital Gyrus |
| | 5 | 101 | .02 | 66.2 | Left Middle Frontal Gyrus |
| | | | | 28.3 | Left Superior Frontal Gyrus |
| | 6 | 166 | .03 | 22.2 | Right Declive |
| | | | | 21.4 | Right Uvula |
| | | | | 16.1 | Right Tuber |
| | 7 | 650 | .012 | 23.8 | Right Cuneus |
| | 8 | 7918 | .003 | 12.9 | Left Inferior Frontal Gyrus |
| | | | | 12.6 | Left Inferior Parietal Lobule |
| | | | | 10.0 | Left Middle Frontal Gyrus |
| | | | | 9.2 | Left Insula |
| | | | | 7.5 | Left Precentral Gyrus |
| | | | | 6.2 | Left Precuneus |
| | | | | 5.6 | Left Postcentral Gyrus |
| | | | | 3.5 | Left Superior Temporal Gyrus |
| | | | | 3.3 | Left Cuneus |

| | | | | | |
|---|---|---|---|---|---|
| | | | | 3.0 | Left Supramarginal Gyrus |
| | | | | 2.0 | Left Superior Parietal Lobule |
| | | | | 1.2 | Left Middle Occipital Gyrus |
| | | | | 1.0 | Left Middle Temporal Gyrus |
| | 9 | 17416 | <.0001 | 16.5 | Right Middle Frontal Gyrus |
| | | | | 10.5 | Right Inferior Frontal Gyrus |
| | | | | 6.7 | Right Superior Frontal Gyrus |
| | | | | 6.7 | Right Medial Frontal Gyrus |
| | | | | 5.6 | Left Medial Frontal Gyrus |
| | | | | 5.0 | Right Cingulate Gyrus |

**Table S3. Cluster information for time-dependent effects of S-ketamine on the phMRI signal for the high S-ketamine dose condition.** Only clusters showing more than 5% overlap with an atlas region are shown. The Daemon (standard Talaraich) atlas was used identification and clusters were extracted using whereami in AFNI.

phMRI: pharmacological magnetic resonance imaging.

| Block | Cluster | # voxels | p-value | Overlap (%) | Region |
|---|---|---|---|---|---|
| 1 | 1 | 6 | .049 | 47.3 | Right Superior Frontal Gyrus |
| | | | | 38.2 | Right Middle Frontal Gyrus |
| | 2 | 16 | .047 | 51.6 | Right Supramarginal Gyrus |
| | | | | 24.5 | Right Angular Gyrus |
| | | | | 23.9 | Right Inferior Parietal Lobule |
| | 3 | 21 | .042 | 18.3 | Right Lingual Gyrus |
| | 4 | 23 | .04 | 25.0 | Right Postcentral Gyrus |
| | 5 | 74 | .038 | 11.3 | Left Superior Frontal Gyrus |
| | 12 | 80 | .033 | 34.8 | Right Superior Parietal Lobule |
| | | | | 19.9 | Right Postcentral Gyrus |
| | | | | 10.2 | Right Precuneus |
| | 13 | 115 | .031 | 47.8 | Left Pyramis |
| | | | | 28.7 | Left Declive |
| | | | | 22.4 | Left Uvula |
| | 14 | 6393 | .001 | 19.6 | Left Middle Frontal Gyrus |
| | | | | 14.6 | Left Insula |
| | | | | 14.3 | Left Inferior Frontal Gyrus |
| | | | | 9.7 | Left Superior Temporal Gyrus |
| | | | | 8.3 | Left Precentral Gyrus |
| | | | | 6.5 | Left Inferior Parietal Lobule |
| | 15 | 18591 | <.0001 | 11.4 | Right Middle Frontal Gyrus |
| | | | | 9.5 | Right Inferior Frontal Gyrus |
| | | | | 6.8 | Right Medial Frontal Gyrus |
| | | | | 6.4 | Right Superior Frontal Gyrus |
| | | | | 5.7 | Left Medial Frontal Gyrus |
| | | | | 5.3 | Left Cingulate Gyrus |
| | | | | 5.3 | Right Cingulate Gyrus |
| 2 | 3 | 38228 | <.0001 | 9.5 | Right Middle Frontal Gyrus |
| | | | | 8.7 | Left Middle Frontal Gyrus |
| | | | | 6.8 | Right Superior Frontal Gyrus |

| | | | | | |
|---|---|---|---|---|---|
| | | | | 5.3 | Right Inferior Frontal Gyrus |
| | | | | 5.0 | Left Superior Frontal Gyrus |
| 5 | 1 | 9 | .046 | 85.1 | Left Medial Frontal Gyrus |
| | | | | 14.9 | Left Cingulate Gyrus |
| | 2 | 10 | .045 | 33.7 | Left Cingulate Gyrus |
| | | | | 30.5 | Right Cingulate Gyrus |
| | | | | 16.8 | Right Paracentral Lobule |
| | 3 | 45 | .041 | 10.5 | Left Paracentral Lobule |
| | | | | 98.7 | Left Cingulate Gyrus |
| | 4 | 65 | .037 | 83.3 | Left Cingulate Gyrus |
| | 5 | 69 | .04 | 68.0 | Right Cingulate Gyrus |
| | | | | 22.6 | Right Medial Frontal Gyrus |
| | 6 | 344 | .015 | 59.5 | Right Insula |
| | | | | 24.8 | Right Inferior Frontal Gyrus |
| | 7 | 1333 | .01 | 24.1 | Right Anterior Cingulate |
| | | | | 21.4 | Left Anterior Cingulate |
| | | | | 20.0 | Right Medial Frontal Gyrus |
| | | | | 16.2 | Left Medial Frontal Gyrus |
| 6 | 1 | 108 | .028 | 47.5 | Right Medial Frontal Gyrus |
| | | | | 16.4 | Left Medial Frontal Gyrus |
| | | | | 8.8 | Right Anterior Cingulate |
| | | | | 7.5 | Right Superior Frontal Gyrus |
| | | | | 5.7 | Left Superior Frontal Gyrus |
| | | | | 5.4 | Left Anterior Cingulate |
| 7 | 1 | 5 | .048 | 56.5 | Right Anterior Cingulate |
| | | | | 13.0 | Right Medial Frontal Gyrus |
| | 2 | 8 | .042 | 93.6 | Right Medial Frontal Gyrus |
| | 3 | 13 | .044 | 97.6 | Left Cingulate Gyrus |
| | 4 | 47 | .038 | 67.0 | Right Inferior Frontal Gyrus |
| | | | | 33.0 | Right Middle Frontal Gyrus |
| | 5 | 52 | .03 | 83.6 | Right Middle Frontal Gyrus |
| | | | | 6.6 | Right Superior Frontal Gyrus |
| | 6 | 115 | .027 | 56.5 | Right Insula |
| | | | | 35.8 | Right Inferior Frontal Gyrus |
| | 7 | 133 | .034 | 45.7 | Right Cingulate Gyrus |
| | | | | 25.9 | Left Cingulate Gyrus |
| | | | | 16.6 | Right Paracentral Lobule |
| | | | | 10.2 | Left Paracentral Lobule |
| 8 | 1 | 33 | .041 | 95.7 | Left Culmen |
| | 2 | 342 | .013 | 46.9 | Right Culmen |
| | | | | 15.0 | Left Culmen |
| | | | | 13.1 | Right Cerebellar Lingual |
| | | | | 8.3 | Right Declive |

**Table S4. Cluster information for dose-dependent effects of S-ketamine on the phMRI signal.** Only clusters consisting of more than 5 voxels and showing more than 5% overlap with an atlas region are shown. The Daemon (standard Talaraich) atlas was used for identification and overlap of the clusters were extracted using whereami in AFNI.

phMRI: pharmacological magnetic resonance imaging.

| **Low dose > placebo** | | | | | |
|---|---|---|---|---|---|
| Block | Cluster | # voxels | p-value | Overlap (%) | Region |
| 1 | 1 | 6 | .044 | 89.3 | Right Medial Frontal Gyrus |
| | 2 | 9 | .041 | 96.5 | Left Insula |
| | 3 | 10 | .044 | 92.7 | Right Anterior Cingulate |
| | 4 | 16 | .035 | 56.8 | Left Superior Temporal Gyrus |
| | | | | 40.6 | Left Transverse Temporal Gyrus |
| | 5 | 85 | .033 | 66.5 | Right Cingulate Gyrus |
| | | | | 16.9 | Left Cingulate Gyrus |
| | 6 | 164 | .026 | 59.8 | Left Medial Frontal Gyrus |
| | | | | 30.2 | Left Anterior Cingulate |
| | 7 | 246 | .019 | 31.1 | Right Medial Frontal Gyrus |
| | | | | 30.7 | Left Medial Frontal Gyrus |
| | | | | 18.5 | Left Anterior Cingulate |
| | | | | 14.6 | Right Anterior Cingulate |
| 2 | 1 | 8 | .038 | 19.2 | Left Postcentral Gyrus |
| | 2 | 33 | .042 | 44.4 | Right Middle Frontal Gyrus |
| | 3 | 56 | .04 | 28.8 | Right Superior Frontal Gyrus |
| | | | | 24.4 | Right Middle Frontal Gyrus |
| | 4 | 82 | .036 | 80.4 | Right Superior Frontal Gyrus |
| | | | | 17.4 | Right Middle Frontal Gyrus |
| | 5 | 1462 | .004 | 44.0 | Right Inferior Frontal Gyrus |
| | | | | 27.4 | Right Middle Frontal Gyrus |
| | | | | 8.6 | Right Insula |
| | 6 | 12834 | <.0001 | 6.8 | Right Precentral Gyrus |
| | | | | 9.4 | Left Middle Frontal Gyrus |
| | | | | 8.5 | Left Medial Frontal Gyrus |
| | | | | 7.2 | Right Medial Frontal Gyrus |
| | | | | 6.4 | Left Superior Frontal Gyrus |
| | | | | 6.2 | Left Inferior Parietal Lobule |
| | | | | 5.5 | Left Precentral Gyrus |
| | | | | 5.2 | Left Precuneus |
| 7 | 1 | 10 | .043 | 100.0 | Left Superior Frontal Gyrus |
| | 2 | 12 | .034 | 100.0 | Left Anterior Cingulate |
| | 3 | 996 | .01 | 22.7 | Left Medial Frontal Gyrus |
| | | | | 20.9 | Right Medial Frontal Gyrus |
| | | | | 16.0 | Right Superior Frontal Gyrus |
| | | | | 13.7 | Right Anterior Cingulate |
| | | | | 10.7 | Right Cingulate Gyrus |
| | | | | 6.0 | Left Anterior Cingulate |

| High dose > placebo | | | | | |
|---|---|---|---|---|---|
| Block | Cluster | # voxels | p-value | Overlap (%) | Region |
| 1 | 1 | 6 | .044 | 5.6 | Left Inferior Frontal Gyrus |
| | 2 | 9 | .049 | 67.0 | Right Middle Frontal Gyrus |
| | 3 | 14 | .037 | 54.1 | Right Insula |
| | | | | 22.2 | Right Claustrum |
| | 4 | 32 | .039 | 92.6 | Right Middle Temporal Gyrus |
| | | | | 6.8 | Right Angular Gyrus |
| | 5 | 132 | .027 | 59.9 | Left Insula |
| | | | | 24.1 | Left Inferior Frontal Gyrus |
| | 6 | 11751 | .004 | 8.6 | Left Middle Frontal Gyrus |
| | | | | 8.0 | Right Middle Frontal Gyrus |
| | | | | 6.9 | Left Medial Frontal Gyrus |
| | | | | 6.7 | Right Superior Frontal Gyrus |
| | | | | 6.6 | Left Superior Frontal Gyrus |
| | | | | 5.7 | Right Inferior Frontal Gyrus |
| | | | | 5.2 | Right Medial Frontal Gyrus |
| | | | | 5.2 | Left Cingulate Gyrus |
| | | | | 5.1 | Left Superior Temporal Gyrus |
| 2 | 1 | 28 | .031 | 19.9 | Right Claustrum |
| | | | | 17.8 | Right Insula |
| | 2 | 21213 | <.0001 | 10.3 | Left Middle Frontal Gyrus |
| | | | | 7.5 | Left Medial Frontal Gyrus |
| | | | | 7.4 | Left Superior Frontal Gyrus |
| | | | | 6.6 | Right Middle Frontal Gyrus |
| | | | | 6.1 | Right Superior Frontal Gyrus |
| | | | | 6.1 | Right Medial Frontal Gyrus |
| | | | | 5.8 | Right Inferior Frontal Gyrus |
| 6 | 1 | 8 | .047 | 70.3 | Left Medial Frontal Gyrus |
| | | | | 20.3 | Left Superior Frontal Gyrus |
| | | | | 9.5 | Left Cingulate Gyrus |
| | 2 | 12 | .048 | 97.3 | Left Superior Frontal Gyrus |
| | 3 | 18 | .04 | 100.0 | Left Middle Frontal Gyrus |
| | 4 | 19 | .043 | 95.7 | Left Middle Frontal Gyrus |
| | 5 | 24 | .044 | 70.2 | Left Precentral Gyrus |
| | | | | 29.4 | Left Middle Frontal Gyrus |
| | 6 | 25 | .038 | 72.3 | Right Medial Frontal Gyrus |
| | | | | 13.9 | Left Medial Frontal Gyrus |
| | 7 | 37 | .032 | 76.4 | Left Medial Frontal Gyrus |
| | | | | 23.6 | Left Superior Frontal Gyrus |
| | 8 | 97 | .023 | 53.8 | Left Middle Frontal Gyrus |
| | | | | 45.0 | Left Superior Frontal Gyrus |
| 7 | 1 | 6 | .047 | 88.5 | Left Middle Frontal Gyrus |
| | 2 | 9 | .043 | 64.4 | Left Middle Frontal Gyrus |
| | | | | 35.6 | Left Precentral Gyrus |

| | | | | |
|---|---|---|---|---|
| 3 | 10 | .041 | 8.3 | Right Middle Frontal Gyrus |
| 4 | 10 | .044 | 100.0 | Left Middle Frontal Gyrus |
| 5 | 11 | .038 | 100.0 | Left Cingulate Gyrus |
| 6 | 12 | .043 | 100.0 | Left Anterior Cingulate |
| 7 | 21 | .04 | 79.4 | Right Inferior Frontal Gyrus |
| 8 | 71 | .041 | 56.2 | Left Middle Frontal Gyrus |
| | | | 23.0 | Left Inferior Frontal Gyrus |
| | | | 20.8 | Left Precentral Gyrus |
| 9 | 690 | .013 | 27.0 | Left Medial Frontal Gyrus |
| | | | 19.3 | Right Medial Frontal Gyrus |
| | | | 13.2 | Left Superior Frontal Gyrus |
| | | | 12.9 | Right Cingulate Gyrus |
| | | | 11.5 | Left Cingulate Gyrus |
| | | | 11.0 | Right Superior Frontal Gyrus |

**Table S5. phMRI response: Linear mixed effect model for individual-level COPE values in the ACC MRS voxel. A.** Bayesian Information Criterion (BIC)=6501. All p-values are corrected using False Discovery Rate (FDR) multiple comparison correction. **B.** Time-dependent effects are reported, i.e. differences between time points within a single condition. **C.** Dose-dependent differences are reported, i.e. differences between conditions per time point. Significant values ($p_{FDR}$<0.05) are highlighted in bold. Model: COPE_fMRI ~ condition * block, random = ~ 1|Subject

ACC: anterior cingulate cortex; COPE: contrast of parameter estimate; MRS; magnetic resonance spectroscopy; phMRI: pharmacological magnetic resonance imaging.

| **A. Linear mixed effects model results** | | |
|---|---|---|
| intercept | F(df)=11.36(1,602) | **p=.0008** |
| condition | F(df)=88.38(2,602) | **p<.0001** |
| block | F(df)=6.10(7,602) | **p<.0001** |
| condition * block | F(df)=1.87(14,602) | **p=.027** |

**B. Post hoc analysis: time-dependent effects**

| | **Block 1 : Block 2** | **Block 1 : Block 3** | **Block 1 : Block 4** | **Block 1 : Block 5** | **Block 1 : Block 6** | **Block 1 : Block 7** | **Block 1 : Block 8** | **Block 2 : Block 3** |
|---|---|---|---|---|---|---|---|---|
| placebo | p=.79 | p=.86 | p=.79 | p=.99 | p=.79 | p=.79 | p=.86 | p=.79 |
| low | p=.97 | **p=.031** | **p=.035** | p=.13 | p=.07 | p=.13 | p=.07 | **p=.032** |
| high | p=.97 | **p=.0001** | **p=.0009** | **p=.005** | **p=.002** | **p=.004** | **p=.004** | **p=.0001** |

| | **Block 2 : Block 4** | **Block 2 : Block 5** | **Block 2 : Block 6** | **Block 2 : Block 7** | **Block 2 : Block 8** | **Block 3 : Block 4** | **Block 3 : Block 5** | **Block 3 : Block 6** |
|---|---|---|---|---|---|---|---|---|
| placebo | p=.86 | p=.79 | p=.86 | p=.86 | p=.86 | p=.86 | p=.86 | p=.86 |
| low | **p=.03** | p=.12 | p=.07 | p=.12 | p=.07 | p=.96 | p=.59 | p=.82 |

| | | | | | | | | |
|---|---|---|---|---|---|---|---|---|
| high | **p=.0008** | **p=.004** | **p=.002** | **p=.004** | **p=.004** | p=.66 | p=.24 | p=.41 |
| | **Block 3 : Block 7** | **Block 3 : Block 8** | **Block 4 : Block 5** | **Block 4 : Block 6** | **Block 4 : Block 7** | **Block 4 : Block 8** | **Block 5 : Block 6** | **Block 5 : Block 7** |
| placebo | p=.86 | p=.89 | p=.79 | p=.99 | p=.98 | p=.86 | p=.79 | p=.79 |
| low | p=.59 | p=.82 | p=.67 | p=.82 | p=.67 | p=.82 | p=.82 | p=.97 |
| high | p=.25 | p=.25 | p=.66 | p=.89 | p=.69 | p=.69 | p=.87 | p=.98 |
| | **Block 5 : Block 8** | **Block 6 : Block 7** | **Block 6 : Block 8** | **Block 7 : Block 8** | | | | |
| placebo | p=.86 | p=.98 | p=.86 | p=.89 | | | | |
| low | p=.82 | p=.82 | p=.97 | p=.82 | | | | |
| high | p=.97 | p=.87 | p=.87 | p=.97 | | | | |

**C. Post hoc analysis: dose-dependent effects**

| | **Block 1** | **Block 2** | **Block 3** | **Block 4** | **Block 5** | **Block 6** | **Block 7** | **Block 8** |
|---|---|---|---|---|---|---|---|---|
| placebo: low | **p<.0001** | **p<.0001** | p=.06 | **p=.005** | **p=.016** | **p=.001** | **p=.0003** | **p=.004** |
| placebo: high | **p<.0001** | **p<.0001** | p=.06 | **p=.0004** | **p=.003** | **p=.0001** | **p<.0001** | **p=.0002** |
| low:high | p=.10 | p=.11 | p=.85 | p=.44 | p=.51 | p=.48 | p=.61 | p=.33 |

**Table S6. phMRI response and heart rate: Linear mixed effect model for individual-level COPE values in the ACC MRS voxel. A.** Bayesian Information Criterium (BIC)=5039 All p-values are corrected using False Discovery Rate (FDR) multiple comparison correction. **B.** Time-dependent effects are reported, i.e. differences between time points within a single condition. **C.** Dose-dependent differences are reported, i.e. differences between conditions per time point. Significant values ($p_{FDR}$<0.05) are highlighted in bold.

Model: COPE_fMRI ~ condition * block + HR_change, random = ~ 1 | Subject

ACC: anterior cingulate cortex; COPE: contrast of parameter estimate; MRS; magnetic resonance spectroscopy; phMRI: pharmacological magnetic resonance imaging.

**A. Linear mixed effects model results**

| | | |
|---|---|---|
| intercept | F(df)=12.11(1,575) | **p=.0005** |
| condition | F(df)=79.73(2, 575) | **p<.0001** |
| block | F(df)=5.99(7, 575) | **p<.0001** |
| %change in heart rate | F(df)=1.69(1,575 | **p=19** |
| condition * block | F(df)=1.83(14, 575) | **p=.032** |

**B. Post hoc analysis: time-dependent effects**

| | Block 1 : Block 2 | Block 1 : Block 3 | Block 1 : Block 4 | Block 1 : Block 5 | Block 1 : Block 6 | Block 1 : Block 7 | Block 1 : Block 8 | Block 2 : Block 3 |
|---|---|---|---|---|---|---|---|---|
| placebo | p=.89 | p=.89 | p=.89 | p=.97 | p=.89 | p=.89 | p=.89 | p=.89 |
| low | p=.98 | **p=.030** | **p=.030** | p=.14 | p=.06 | p=.08 | **p=.030** | **p=.030** |
| high | p=.89 | **p=.0004** | **p=.002** | **p=.007** | **p=.003** | **p=.005** | **p=.005** | **p=.0001** |
| | **Block 2 : Block 4** | **Block 2 : Block 5** | **Block 2 : Block 6** | **Block 2 : Block 7** | **Block 2 : Block 8** | **Block 3 : Block 4** | **Block 3 : Block 5** | **Block 3 : Block 6** |
| placebo | p=.89 | p=.89 | p=.89 | p=.89 | p=.89 | p=.89 | p=.89 | p=.89 |
| low | **p=.030** | p=.10 | **p=.044** | p=.06 | **p=.030** | p=.99 | p=.71 | p=.88 |
| high | **p=.0008** | **p=.004** | **p=.002** | **p=.003** | **p=.003** | p=.75 | p=.36 | p=.59 |
| | **Block 3 : Block 7** | **Block 3 : Block 8** | **Block 4 : Block 5** | **Block 4 : Block 6** | **Block 4 : Block 7** | **Block 4 : Block 8** | **Block 5 : Block 6** | **Block 5 : Block 7** |
| placebo | p=.89 | p=.89 | p=.89 | p=.97 | p=.97 | p=.96 | p=.89 | p=.89 |
| low | p=.88 | p=.99 | p=.71 | p=.88 | p=.88 | p=.99 | p=.88 | p=.88 |
| high | p=.41 | p=.41 | p=.75 | p=.89 | p=.81 | p=.81 | p=.89 | p=.92 |
| | **Block 5 : Block 8** | **Block 6 : Block 7** | **Block 6 : Block 8** | **Block 7 : Block 8** | | | | |
| placebo | p=.89 | p=.97 | p=.97 | p=.97 | | | | |
| low | p=.71 | p=.99 | p=.88 | p=.88 | | | | |
| high | p=.92 | p=.89 | p=.89 | p>.99 | | | | |

**C. Post hoc analysis: dose-dependent effects**

| | Block 1 | Block 2 | Block 3 | Block 4 | Block 5 | Block 6 | Block 7 | Block 8 |
|---|---|---|---|---|---|---|---|---|
| placebo: low | **p<.0001** | **p<.0001** | p=.06 | **p=.009** | **p=.017** | **p=.003** | **p=.002** | **p=.020** |
| placebo: high | **p<.0001** | **p<.0001** | p=.06 | **p=.0006** | **p=.004** | **p=.0003** | **p=.0001** | **p=.0002** |
| low:high | p=.16 | p=.15 | p=.88 | p=.39 | p=.58 | p=.51 | p=.43 | p=.19 |

**Table S7. phMRI response and framewise displacement (FD): Linear mixed effect model for individual-level COPE values in the ACC MRS voxel. A.** Bayesian Information Criterium (BIC)=6577. All p-values are corrected using False Discovery Rate (FDR) multiple comparison correction. **B.** Time-dependent effects are reported, i.e. differences between time points within a single condition. **C.** Dose-dependent differences are reported, i.e. differences between conditions per time point. Significant values ($p_{FDR}$<0.05) are highlighted in bold.

Model: COPE_fMRI ~ condition * block + FD * condition, random = ~ 1 | Subject

ACC: anterior cingulate cortex; COPE: contrast of parameter estimate; MRS; magnetic resonance spectroscopy; phMRI: pharmacological magnetic resonance imaging.

| **A. Linear mixed effects model results** | | |
|---|---|---|
| intercept | F(df)=7.81(1,578) | **p=.0054** |
| condition | F(df)=97.40(2,578) | **p<.0001** |
| block | F(df)=6.78(7,578) | **p<.0001** |
| FD | F(df)=17.57(1,578) | **p<.0001** |
| condition * block | F(df)=2.07(14,578) | **p=.012** |
| condition * FD | F(df)=16.67(2,578) | **p<.0001** |
| block * FD | F(df)=0.75(7,578) | **p=.63** |
| condition * block * FD | F(df)=1.79(14,578) | **p=.037** |

**B. Post hoc analysis: time-dependent effects**

| | **Block 1 : Block 2** | **Block 1 : Block 3** | **Block 1 : Block 4** | **Block 1 : Block 5** | **Block 1 : Block 6** | **Block 1 : Block 7** | **Block 1 : Block 8** | **Block 2 : Block 3** |
|---|---|---|---|---|---|---|---|---|
| placebo | p=.84 | p=.89 | p=.84 | p=.97 | p=.84 | p=.84 | p=.88 | p=.84 |
| low | p=.78 | **p=.018** | **p=.027** | p=.14 | p=.06 | p=.15 | p=.06 | **p=.009** |
| high | p=.94 | **p<.0001** | **p=.0004** | **p=.004** | **p=.002** | **p=.003** | **p=.003** | **p<.0001** |
| | **Block 2 : Block 4** | **Block 2 : Block 5** | **Block 2 : Block 6** | **Block 2 : Block 7** | **Block 2 : Block 8** | **Block 3 : Block 4** | **Block 3 : Block 5** | **Block 3 : Block 6** |
| placebo | p=.88 | p=.84 | p=.89 | p=.88 | p=.88 | p=.88 | p=.88 | p=.88 |
| low | **p=.009** | **p=.039** | **p=.018** | **p=.042** | **p=.018** | p=.91 | p=.61 | p=.78 |
| high | **p=.0002** | **p=.003** | **p=.0008** | **p=.002** | **p=.002** | p=.60 | p=.19 | p=.37 |
| | **Block 3 : Block 7** | **Block 3 : Block 8** | **Block 4 : Block 5** | **Block 4 : Block 6** | **Block 4 : Block 7** | **Block 4 : Block 8** | **Block 5 : Block 6** | **Block 5 : Block 7** |
| placebo | p=.88 | p=.89 | p=.84 | p=.97 | p=.97 | p=.89 | p=.84 | p=.84 |
| low | p=.58 | p=.78 | p=.71 | p=.78 | p=.68 | p=.78 | p=.78 | p=.96 |
| high | p=.20 | p=.20 | p=.60 | p=.83 | p=.60 | p=.60 | p=.83 | p=.97 |

| | Block 5 : Block 8 | Block 6 : Block 7 | Block 6 : Block 8 | Block 7 : Block 8 |
|---|---|---|---|---|
| placebo | p=.88 | p=.97 | p=.89 | p=.89 |
| low | p=.78 | p=.78 | p=.96 | p=.78 |
| high | p=.97 | p=.83 | p=.83 | p=.99 |

**C. Post hoc analysis: dose-dependent effects**

| | Block 1 | Block 2 | Block 3 | Block 4 | Block 5 | Block 6 | Block 7 | Block 8 |
|---|---|---|---|---|---|---|---|---|
| placebo:low | **p<.0001** | **p<.0001** | **p=.025** | **p=.009** | **p=.004** | **p=.0002** | **p<.0001** | **p=.0007** |
| placebo:high | **p<.0001** | **p<.0001** | **p=.025** | **p=.0003** | **p=.002** | **p<.0001** | **p<.0001** | **p=.0001** |
| low:high | p=.18 | p=.39 | p=.78 | p=.76 | p=.76 | p=.75 | p=.89 | p=.53 |

**Table S8. phMRI response and medication order: Linear mixed effect model for individual-level COPE values in the ACC MRS voxel**. **A.** Bayesian Information Criterium (BIC)=6372. All p-values are corrected using False Discovery Rate (FDR) multiple comparison correction. **B.** Time-dependent effects are reported, i.e. differences between time points within a single condition. **C.** Dose-dependent differences are reported, i.e. differences between conditions per time point. Significant values ($p_{FDR}$<0.05) are highlighted in bold.

Model: COPE_fMRI ~ condition * block + order_med

ACC: anterior cingulate cortex; COPE: contrast of parameter estimate; MRS; magnetic resonance spectroscopy; phMRI: pharmacological magnetic resonance imaging.

**A. Linear mixed effects model results**

| | | |
|---|---|---|
| intercept | F(df)=11.08(1,602) | **p=.0009** |
| condition | F(df)=88.32(2, 602) | **p<.0001** |
| block | F(df)=6.10(7, 602) | **p<.0001** |
| medication order | F(df)=0.15(1,29) | p=.71 |
| condition * block | F(df)=1.87(14, 602) | **p=.027** |

**B. Post hoc analysis: time-dependent effects**

| | Block 1 : Block 2 | Block 1 : Block 3 | Block 1 : Block 4 | Block 1 : Block 5 | Block 1 : Block 6 | Block 1 : Block 7 | Block 1 : Block 8 | Block 2 : Block 3 |
|---|---|---|---|---|---|---|---|---|
| placebo | p=.79 | p=.86 | p=.79 | p=.99 | p=.79 | p=.79 | p=.86 | p=.79 |
| low | p=.97 | **p=.032** | **p=.035** | p=.13 | p=.07 | p=.13 | p=.07 | **p=.032** |
| high | p=.97 | **p=.0001** | **p=.0009** | **p=.005** | **p=.002** | **p=.004** | **p=.004** | **p=.0001** |

| | Block 2 : Block 4 | Block 2 : Block 5 | Block 2 : Block 6 | Block 2 : Block 7 | Block 2 : Block 8 | Block 3 : Block 4 | Block 3 : Block 5 | Block 3 : Block 6 |
|---|---|---|---|---|---|---|---|---|
| placebo | p=.86 | p=.79 | p=.86 | p=.86 | p=.86 | p=.86 | p=.86 | p=.86 |
| low | **p=.032** | p=.12 | p=.07 | p=.12 | p=.07 | p=.97 | p=.59 | p=.82 |
| high | **p=.0008** | **p=.004** | **p=.002** | **p=.004** | **p=.004** | p=.66 | p=.24 | p=.41 |
| | **Block 3 : Block 7** | **Block 3 : Block 8** | **Block 4 : Block 5** | **Block 4 : Block 6** | **Block 4 : Block 7** | **Block 4 : Block 8** | **Block 5 : Block 6** | **Block 5 : Block 7** |
| placebo | p=.86 | p=.89 | p=.79 | p=.99 | p=.98 | p=.86 | p=.79 | p=.79 |
| low | p=.59 | p=.82 | p=.67 | p=.82 | p=.67 | p=.82 | p=.82 | p=.97 |
| high | p=.25 | p=.25 | p=.66 | p=.89 | p=.69 | p=.69 | p=.89 | p=.97 |
| | **Block 5 : Block 8** | **Block 6 : Block 7** | **Block 6 : Block 8** | **Block 7 : Block 8** | | | | |
| placebo | p=.86 | p=.98 | p=.86 | p=.89 | | | | |
| low | p=.82 | p=.82 | p=.97 | p=.82 | | | | |
| high | p=.97 | p=.89 | p=.89 | p=.99 | | | | |

**C. Post hoc analysis: dose-dependent effects**

| | Block 1 | Block 2 | Block 3 | Block 4 | Block 5 | Block 6 | Block 7 | Block 8 |
|---|---|---|---|---|---|---|---|---|
| placebo: low | **p<.0001** | **p<.0001** | p=.06 | **p=.005** | **p=.016** | **p=.001** | **p=.0003** | **p=.004** |
| placebo: high | **p<.0001** | **p<.0001** | p=.06 | **p=.0004** | **p=.003** | **p=.0001** | **p<.0001** | **p=.0002** |
| low:high | p=.10 | p=.11 | p=.85 | p=.43 | p=.50 | p=.47 | p=.60 | p=.33 |

**Table S9. Heart rate: linear mixed effect model for %change in heart rate from baseline. A.** Bayesian Information Criterium (BIC)=5039. All p-values are corrected using False Discovery Rate (FDR) multiple comparison correction. **B.** Time-dependent effects are reported, i.e. differences between time points within a single condition. **C.** Dose-dependent differences are reported, i.e. differences between conditions per time point. Significant values ($p_{FDR}$<0.05) are highlighted in bold.
Model: HR_change ~ condition * block, random = ~ 1 | Subject

| **A. Linear mixed effects model results** | | |
|---|---|---|
| intercept | F(df)=10.94(1,595) | **p=.001** |
| condition | F(df)=32.07(2,595) | **p<.0001** |
| block | F(df)=8.30(7,595) | **p<.0001** |
| condition * block | F(df)=2.51(14,595) | **p=.002** |

| **B. Post hoc analysis: time-dependent effects** | | | | | | | | |
|---|---|---|---|---|---|---|---|---|
| | **Block 1 : Block 2** | **Block 1 : Block 3** | **Block 1 : Block 4** | **Block 1 : Block 5** | **Block 1 : Block 6** | **Block 1 : Block 7** | **Block 1 : Block 8** | **Block 2 : Block 3** |
| placebo | p=.95 | p=.95 | p=.95 | p=.95 | p=.98 | p=.95 | p=.95 | p=.95 |
| low | p=.15 | p=.26 | p=.92 | p=.74 | p=.25 | p=.15 | p=.15 | p=.74 |
| high | **p=.001** | **p<.0001** | **p=.001** | p=.09 | p=.16 | p=.38 | p=.79 | **p=.035** |
| | **Block 2 : Block 4** | **Block 2 : Block 5** | **Block 2 : Block 6** | **Block 2 : Block 7** | **Block 2 : Block 8** | **Block 3 : Block 4** | **Block 3 : Block 5** | **Block 3 : Block 6** |
| placebo | p=.95 | p=.95 | p=.95 | p=.95 | p=.95 | p=.98 | p=.95 | p=.95 |
| low | p=.15 | p=.07 | **p=.008** | **p=.003** | **p=.003** | p=.25 | p=.15 | p=.024 |
| high | p=.93 | p=.12 | p=.06 | **p=.020** | **p=.002** | **p=.028** | **p=.0001** | **p<.0001** |
| | **Block 3 : Block 7** | **Block 3 : Block 8** | **Block 4 : Block 5** | **Block 4 : Block 6** | **Block 4 : Block 7** | **Block 4 : Block 8** | **Block 5 : Block 6** | **Block 5 : Block 7** |
| placebo | p=.98 | p=.95 | p=.95 | p=.95 | p=.98 | p=.95 | p=.95 | p=.95 |
| low | **p=.008** | **p=.008** | p=.76 | p=.26 | p=.16 | p=.16 | p=.45 | p=.26 |
| high | **p<.0001** | **p<.0001** | p=.13 | p=.07 | **p=.022** | **p=.003** | p=.75 | p=.43 |
| | **Block 5 : Block 8** | **Block 6 : Block 7** | **Block 6 : Block 8** | **Block 7 : Block 8** | | | | |
| placebo | p=.98 | p=.96 | p=.95 | p=.95 | | | | |
| low | p=.26 | p=.75 | p=.75 | p=.97 | | | | |
| high | p=.14 | p=.66 | p=.27 | p=.54 | | | | |

| **C. Post hoc analysis: dose-dependent effects** | | | | | | | | |
|---|---|---|---|---|---|---|---|---|
| | **Block 1** | **Block 2** | **Block 3** | **Block 4** | **Block 5** | **Block 6** | **Block 7** | **Block 8** |
| placebo: low | p=.98 | p=.26 | p=.09 | p=.82 | p=.65 | p=.14 | p=.15 | p=.25 |
| placebo: high | p=.98 | **p=.012** | **p<.0001** | **p=.0002** | **p=.009** | p=.14 | p=.18 | p=.25 |
| low:high | p=.98 | p=.14 | **p<.0001** | **p=.0004** | **p=.022** | **p=.007** | **p=.009** | p=.06 |

**Table S10. Cluster information for time-dependent effects of S-ketamine on the phMRI signal using a signal model analysis.** Only clusters consisting of more than 5 voxels and showing more than 5% overlap with an atlas region are shown. The Daemon (standard Talaraich) atlas was used for identification and overlap of the clusters were extracted using whereami in AFNI.
phMRI: pharmacological magnetic resonance imaging.

| Cluster | # voxels | p-value | Overlap (%) | Region |
|---|---|---|---|---|
| **Low S-ketamine dose** | | | | |
| 1 | 50369 | <.0001 | 8.1 | Right Middle Frontal Gyrus |
| | | | 6.6 | Left Middle Frontal Gyrus |
| | | | 5.6 | Right Superior Frontal Gyrus |
| **High S-ketamine dose** | | | | |
| 1 | 87346 | <.0001 | 5.3 | Right Middle Frontal Gyrus |
| | | | 5.0 | Left Middle Frontal Gyrus |

**Table S11. Cluster information for dose-dependent effects of S-ketamine on the phMRI signal using a signal model analysis.** Only clusters consisting of more than 5 voxels and showing more than 5% overlap with an atlas region are shown. The Daemon (standard Talaraich) atlas was used for identification and overlap of the clusters were extracted using whereami in AFNI.
phMRI: pharmacological magnetic resonance imaging.

| Cluster | # voxels | p-value | Overlap (%) | Region |
|---|---|---|---|---|
| **Placebo vs low S-ketamine dose** | | | | |
| 1 | 11 | 0.047 | 100 | Right Superior Temporal Gyrus |
| 2 | 14 | 0.038 | 54.7 | Right Superior Temporal Gyrus |
| 3 | 35579 | <.0001 | 7.4 | Right Middle Frontal Gyrus |
| | | | 6.9 | Left Middle Frontal Gyrus |
| **Placebo vs high S-ketamine dose** | | | | |
| 1 | 31 | 0.039 | 30 | Left Culmen |
| 2 | 1119 | 0.013 | 16 | Left Declive |
| | | | 13.5 | Right Thalamus |
| | | | 13.2 | Left Thalamus |
| | | | 11.5 | Left Culmen |
| | | | 5.4 | Left Parahippocampal Gyrus |
| | | | 5.3 | Left Uvula |
| 3 | 53939 | <.0001 | 6.6 | Left Middle Frontal Gyrus |
| | | | 6.5 | Right Middle Frontal Gyrus |

**Table S12. Cluster information for differences in the phMRI signal in the placebo condition for subjects who had prior experience with ketamine and those without.** Only clusters consisting of more than 5 voxels and showing more than 5% overlap with an atlas region are shown. The Daemon (standard Talaraich) atlas was used for identification and overlap of the clusters were extracted using whereami in AFNI.

phMRI: pharmacological magnetic resonance imaging

| Block | Cluster | # voxels | p-value | Overlap (%) | Region |
|---|---|---|---|---|---|
| 2 | 1 | 5 | .046 | 100.0% | Left Precuneus |
| | 2 | 5 | .048 | 33.3 | Right Insula |
| | 3 | 6 | .043 | 96.4 | Left Postcentral Gyrus |
| | 4 | 6 | .048 | 100.0% | Right Postcentral Gyrus |
| | 5 | 7 | .046 | 56.9 | Left Cuneus |
| | | | | 20.0 | Left Precuneus |
| | 6 | 8 | .047 | 52.7 | Right Precentral Gyrus |
| | | | | 47.3 | Right Postcentral Gyrus |
| | 7 | 8 | .047 | 73.2 | Right Postcentral Gyrus |
| | | | | 26.8 | Right Inferior Parietal Lobule |
| | 8 | 9 | .046 | 100.0% | Right Precuneus |
| | 9 | 11 | .045 | 55.2 | Left Cuneus |
| | | | | 43.8 | Left Precuneus |
| | 10 | 29 | .042 | 99.3 | Right Cuneus |
| | 11 | 62 | .036 | 89.2 | Right Precentral Gyrus |
| | | | | 10.8 | Right Postcentral Gyrus |
| | 12 | 112 | .03 | 47.7 | Left Cuneus |
| | | | | 43.8 | Left Lingual Gyrus |
| 7 | 1 | 3 | .043 | 12.5 | Right Postcentral Gyrus |
| | 2 | 7 | .046 | 100.0% | Right Inferior Parietal Lobule |
| | 3 | 8 | .044 | 40.0 | Left Cuneus |
| | | | | 21.3 | Left Precuneus |
| | | | | 12.0 | Right Precuneus |
| | 4 | 41 | .043 | 52.8 | Right Cuneus |
| | 5 | 54 | .035 | 31.0 | Right Cuneus |
| | | | | 12.2 | Left Cuneus |
| | 6 | 159 | .018 | 63.2 | Right Precentral Gyrus |
| | | | | 33.2 | Right Postcentral Gyrus |
| | 7 | 1708 | .015 | 29.8 | Left Precuneus |
| | | | | 17.8 | Right Precuneus |
| | | | | 14.6 | Left Cuneus |

**Table S13. phMRS: Time-dependent effects of S-ketamine on neurometabolite levels in the ACC.** Per condition (placebo, low dose of S-ketamine, and high dose of S-ketamine), and per infusion block, the % change from baseline with standard deviation (SD) are provided, as well as the original p-values and the false discovery rate (FDR)-corrected p-values ($p_{FDR}$).

ACC: anterior cingulate cortex; phMRS: pharmacological magnetic resonance spectroscopy.

**Placebo**

| | Block 1 | | | | Block 2 | | | | Block 3 | | | | Block 4 | | | |
|---|---|---|---|---|---|---|---|---|---|---|---|---|---|---|---|---|
| | % | ±SD | p | $p_{FDR}$ | % | ±SD | p | $p_{FDR}$ | % | ±SD | p | $p_{FDR}$ | % | ±SD | p | $p_{FDR}$ |
| Asc | -0.27 | 4.38 | 0.48 | 0.480 | 5.21 | 4.61 | **0.13** | 0.360 | 3.77 | 6.03 | **0.27** | 0.390 | 2.03 | 5.01 | 0.34 | 0.390 |
| Asp | -0.33 | 1.98 | 0.44 | 0.440 | 1.74 | 1.82 | 0.17 | 0.370 | 0.55 | 2.13 | 0.40 | 0.440 | 0.53 | 2.24 | 0.41 | 0.440 |
| tCr | -0.25 | 0.32 | 0.22 | 0.250 | 0.34 | 0.51 | 0.26 | 0.260 | -0.45 | 0.52 | 0.20 | 0.250 | -0.64 | 0.64 | 0.17 | 0.250 |
| GSH | 0.19 | 1.36 | 0.44 | 0.440 | 4.11 | 1.34 | **0.004** | **0.010** | 2.61 | 1.57 | 0.05 | 0.070 | 3.83 | 1.31 | **0.004** | **0.010** |
| Glu | -0.70 | 0.64 | 0.14 | 0.170 | 1.13 | 0.99 | 0.13 | 0.170 | -0.92 | 0.82 | 0.13 | 0.170 | -0.64 | 0.78 | 0.21 | 0.210 |
| Ins | 0.13 | 0.36 | 0.36 | 0.410 | 0.19 | 0.42 | 0.33 | 0.410 | -0.37 | 0.76 | 0.31 | 0.410 | -0.32 | 0.49 | 0.26 | 0.410 |
| Lac | 3.95 | 3.36 | 0.13 | 0.270 | 4.80 | 4.24 | 0.13 | 0.270 | 8.14 | 4.97 | 0.06 | 0.270 | 3.98 | 4.07 | 0.17 | 0.270 |
| tNAA | -0.13 | 0.24 | 0.29 | 0.290 | -0.18 | 0.24 | 0.23 | 0.290 | -0.16 | 0.26 | 0.28 | 0.290 | -0.43 | 0.39 | 0.14 | 0.220 |
| tCh | 0.66 | 0.68 | 0.17 | 0.370 | 0.78 | 0.72 | 0.14 | 0.370 | 0.86 | 0.75 | 0.13 | 0.370 | 0.47 | 0.79 | 0.28 | 0.370 |
| PE | -2.02 | 1.82 | 0.14 | 0.140 | -4.08 | 1.28 | **0.002** | **0.002** | -6.17 | 1.79 | **0.001** | **0.002** | -6.18 | 1.46 | **0.0001** | **0.0004** |
| Scyllo | -2.18 | 2.90 | 0.23 | 0.380 | -0.84 | 2.92 | 0.39 | 0.390 | -2.96 | 3.18 | 0.18 | 0.390 | -6.35 | 2.92 | **0.02** | 0.160 |
| Glc+Tau | -0.50 | 1.04 | 0.32 | 0.320 | -3.06 | 1.36 | **0.02** | 0.060 | -2.24 | 1.54 | 0.08 | 0.090 | -1.94 | 1.22 | 0.06 | 0.080 |
| sigma | 0.38 | 0.97 | 0.35 | 0.350 | 3.12 | 1.73 | **0.04** | 0.170 | 2.64 | 1.96 | 0.10 | 0.250 | 3.47 | 1.71 | **0.03** | 0.170 |

| | Block 5 | | | | Block 6 | | | | Block 7 | | | | Block 8 | | | |
|---|---|---|---|---|---|---|---|---|---|---|---|---|---|---|---|---|
| | % | ±SD | p | $p_{FDR}$ | % | ±SD | p | $p_{FDR}$ | % | ±SD | p | $p_{FDR}$ | % | ±SD | p | $p_{FDR}$ |
| Asc | 3.00 | 5.60 | **0.30** | 0.390 | 8.01 | 4.59 | **0.05** | 0.360 | 6.32 | 4.98 | **0.11** | 0.360 | 3.28 | 4.47 | 0.24 | 0.390 |
| Asp | 0.45 | 2.55 | 0.43 | 0.440 | 3.21 | 1.99 | 0.06 | 0.370 | 2.21 | 2.18 | 0.16 | 0.370 | 1.90 | 2.06 | 0.18 | 0.370 |
| tCr | -1.00 | 0.69 | 0.08 | 0.190 | -0.88 | 0.65 | 0.09 | 0.190 | -0.91 | 0.58 | 0.06 | 0.190 | -1.91 | 0.67 | **0.00** | **0.030** |
| GSH | 3.61 | 1.17 | **0.00** | **0.010** | 2.83 | 1.37 | **0.02** | 0.050 | 1.95 | 1.83 | 0.15 | 0.170 | 2.62 | 1.47 | **0.04** | 0.070 |
| Glu | -2.12 | 1.09 | **0.03** | 0.060 | -2.06 | 0.83 | **0.01** | **0.030** | -2.46 | 0.89 | **0.01** | **0.030** | -2.72 | 1.01 | **0.01** | **0.030** |
| Ins | -0.32 | 0.59 | 0.29 | 0.410 | -0.03 | 0.58 | 0.48 | 0.480 | -0.49 | 0.49 | 0.17 | 0.410 | -0.92 | 0.73 | 0.11 | 0.410 |
| Lac | -0.99 | 4.10 | 0.41 | 0.590 | 3.26 | 3.36 | 0.17 | 0.270 | -1.00 | 3.78 | 0.40 | 0.490 | -0.93 | 5.18 | 0.43 | 0.490 |
| tNAA | -0.71 | 0.37 | **0.03** | 0.090 | -0.67 | 0.41 | 0.06 | 0.110 | -1.19 | 0.49 | **0.01** | **0.040** | -1.18 | 0.48 | **0.01** | **0.040** |

| | | | | | | | | | | | | | | | | |
|---|---|---|---|---|---|---|---|---|---|---|---|---|---|---|---|---|
| tCh | -0.07 | 0.96 | 0.47 | 0.470 | 0.69 | 0.98 | 0.24 | 0.372 | 0.73 | 0.98 | 0.23 | 0.300 | 0.20 | 0.94 | 0.42 | 0.470 |
| PE | -5.77 | 1.51 | **.0004** | **0.001** | -6.73 | 2.10 | **0.002** | **0.002** | -9.27 | 1.29 | **<.0001** | **<.0001** | -9.34 | 1.62 | **<.0001** | **<.0001** |
| Scyllo | -1.15 | 2.92 | 0.35 | 0.390 | -1.96 | 2.92 | 0.25 | 0.390 | -1.23 | 2.93 | 0.34 | 0.390 | -4.94 | 2.93 | 0.05 | 0.210 |
| Glc+Tau | -2.88 | 1.40 | **0.03** | 0.060 | -2.84 | 1.46 | **0.03** | 0.060 | -3.95 | 1.38 | **0.004** | **0.030** | -2.58 | 1.49 | **0.05** | 0.070 |
| sigma | 1.90 | 1.90 | 0.16 | 0.290 | 1.88 | 2.04 | 0.18 | 0.290 | 1.21 | 2.19 | 0.29 | 0.330 | 1.42 | 2.55 | 0.29 | 0.330 |
| **Low dose** | | | | | | | | | | | | | | | | |
| | Block 1 | | | | Block 2 | | | | Block 3 | | | | Block 4 | | | |
| | % | ±SD | p | $p_{FDR}$ | % | ±SD | p | $p_{FDR}$ | % | ±SD | p | $p_{FDR}$ | % | ±SD | p | $p_{FDR}$ |
| Asc | 1.49 | 3.72 | **0.35** | 0.420 | -1.14 | 3.64 | **0.38** | 0.420 | 0.93 | 3.34 | **0.39** | 0.420 | -3.03 | 3.39 | 0.19 | 0.420 |
| Asp | 3.89 | 1.86 | **0.02** | 0.190 | -1.48 | 1.86 | 0.22 | 0.290 | 0.34 | 2.26 | 0.44 | 0.440 | 2.24 | 2.16 | 0.16 | 0.250 |
| tCr | 0.18 | 0.38 | 0.32 | 0.430 | 1.20 | 0.41 | **0.004** | **0.030** | 1.06 | 0.50 | **0.02** | 0.090 | 0.48 | 0.46 | 0.15 | 0.240 |
| GSH | 1.63 | 1.74 | 0.18 | 0.250 | 1.46 | 1.82 | 0.22 | 0.250 | 1.89 | 1.94 | 0.17 | 0.250 | 1.17 | 1.71 | 0.25 | 0.250 |
| Glu | 0.67 | 0.65 | 0.16 | 0.320 | 2.21 | 1.02 | **0.02** | 0.150 | 1.60 | 0.86 | **0.04** | 0.150 | 0.14 | 0.86 | 0.43 | 0.490 |
| Ins | 0.28 | 0.68 | 0.34 | 0.430 | 0.78 | 0.71 | 0.14 | 0.430 | 0.20 | 0.78 | 0.40 | 0.430 | 0.47 | 0.75 | 0.27 | 0.430 |
| Lac | 3.69 | 4.52 | 0.21 | 0.420 | 12.77 | 3.50 | **.0006** | **0.005** | 9.24 | 3.80 | **0.01** | **0.046** | 4.98 | 3.33 | 0.07 | 0.200 |
| tNAA | 0.19 | 0.21 | 0.18 | 0.210 | 0.63 | 0.31 | **0.03** | 0.110 | 0.46 | 0.19 | **0.01** | 0.100 | 0.36 | 0.24 | 0.07 | 0.150 |
| tCh | 0.11 | 0.77 | 0.44 | 0.440 | 0.46 | 0.65 | 0.24 | 0.320 | 1.44 | 0.69 | **0.02** | 0.070 | 0.37 | 0.81 | 0.33 | 0.370 |
| PE | 0.43 | 2.69 | 0.44 | 0.440 | 2.70 | 2.36 | 0.13 | 0.210 | -2.10 | 2.25 | 0.18 | 0.210 | 2.09 | 2.21 | 0.18 | 0.210 |
| Scyllo | 0.43 | 2.83 | 0.44 | 0.440 | -1.41 | 3.51 | 0.35 | 0.440 | -3.77 | 3.10 | 0.12 | 0.440 | -0.79 | 2.87 | 0.39 | 0.440 |
| Glc+Tau | -1.67 | 1.34 | 0.11 | 0.290 | -1.37 | 1.39 | 0.17 | 0.290 | -1.83 | 1.42 | 0.10 | 0.290 | -0.78 | 1.26 | 0.27 | 0.310 |
| sigma | -3.12 | 1.00 | **0.002** | **0.019** | -2.59 | 1.69 | 0.07 | 0.140 | -0.35 | 1.87 | 0.43 | 0.500 | -0.23 | 1.58 | 0.44 | 0.500 |
| | Block 5 | | | | Block 6 | | | | Block 7 | | | | Block 8 | | | |
| | % | ±SD | p | $p_{FDR}$ | % | ±SD | p | $p_{FDR}$ | % | ±SD | p | $p_{FDR}$ | % | ±SD | p | $p_{FDR}$ |
| Asc | 2.29 | 2.87 | **0.22** | 0.420 | -3.83 | 3.70 | **0.16** | 0.420 | -0.79 | 3.74 | **0.42** | 0.420 | 3.23 | 4.30 | 0.23 | 0.420 |
| Asp | 2.07 | 1.91 | 0.14 | 0.250 | 3.51 | 2.46 | 0.08 | 0.250 | 0.76 | 1.90 | 0.35 | 0.400 | 3.28 | 2.58 | 0.11 | 0.250 |
| tCr | 0.92 | 0.52 | **0.05** | 0.120 | 0.60 | 0.49 | 0.12 | 0.230 | 0.19 | 0.67 | 0.39 | 0.430 | -0.11 | 0.62 | 0.43 | 0.430 |
| GSH | 1.16 | 1.62 | 0.24 | 0.250 | 5.15 | 2.41 | **0.02** | 0.170 | 1.23 | 1.78 | 0.25 | 0.250 | 1.48 | 1.56 | 0.18 | 0.250 |
| Glu | -0.02 | 0.82 | 0.49 | 0.490 | -0.08 | 0.85 | 0.46 | 0.490 | -0.57 | 1.09 | 0.30 | 0.480 | -1.20 | 1.03 | 0.13 | 0.320 |
| Ins | 0.88 | 0.69 | 0.11 | 0.430 | 0.16 | 0.88 | 0.43 | 0.430 | 0.39 | 0.82 | 0.32 | 0.430 | -0.16 | 0.91 | 0.43 | 0.430 |
| Lac | 1.46 | 4.09 | 0.36 | 0.440 | -2.20 | 4.05 | 0.30 | 0.440 | 0.85 | 3.50 | 0.41 | 0.440 | -0.54 | 3.43 | 0.44 | 0.440 |
| tNAA | 0.35 | 0.24 | 0.08 | 0.150 | 0.11 | 0.28 | 0.34 | 0.340 | -0.40 | 0.30 | 0.10 | 0.160 | -0.35 | 0.37 | 0.17 | 0.210 |

| | | | | | | | | | | | | | | | | |
|---|---|---|---|---|---|---|---|---|---|---|---|---|---|---|---|---|
| tCh | 1.86 | 0.85 | **0.02** | 0.070 | 1.69 | 0.82 | **0.03** | 0.070 | 1.24 | 1.03 | 0.12 | 0.240 | 1.04 | 1.01 | 0.16 | 0.250 |
| PE | -3.93 | 2.73 | 0.08 | 0.210 | -2.76 | 2.11 | 0.10 | 0.210 | -3.14 | 2.86 | 0.14 | 0.210 | -4.89 | 2.88 | 0.05 | 0.210 |
| Scyllo | -1.89 | 2.88 | 0.26 | 0.440 | 0.57 | 3.41 | 0.43 | 0.440 | -2.50 | 3.46 | 0.24 | 0.440 | -5.37 | 3.58 | 0.07 | 0.440 |
| Glc+Tau | -1.07 | 1.36 | 0.22 | 0.290 | -1.01 | 1.04 | 0.17 | 0.290 | -0.27 | 1.34 | 0.42 | 0.420 | -1.26 | 1.58 | 0.22 | 0.290 |
| sigma | -0.02 | 1.69 | 0.49 | 0.500 | -1.79 | 1.79 | 0.16 | 0.260 | -3.60 | 2.07 | **0.048** | 0.130 | -3.94 | 2.16 | **0.04** | 0.130 |
| **High dose** | | | | | | | | | | | | | | | | |
| | Block 1 | | | | Block 2 | | | | Block 3 | | | | Block 4 | | | |
| | % | ±SD | p | $p_{FDR}$ | % | ±SD | p | $p_{FDR}$ | % | ±SD | p | $p_{FDR}$ | % | ±SD | p | $p_{FDR}$ |
| Asc | 4.71 | 3.35 | **0.086** | 0.280 | 2.37 | 3.52 | **0.254** | 0.290 | 3.67 | 3.36 | **0.142** | 0.280 | 3.88 | 3.38 | 0.131 | 0.280 |
| Asp | -3.67 | 1.51 | **0.011** | 0.088 | -3.22 | 1.75 | **0.038** | 0.120 | -2.52 | 1.58 | 0.062 | 0.120 | -0.81 | 1.43 | 0.288 | 0.330 |
| tCr | 0.82 | 0.38 | **0.021** | **0.021** | 2.60 | 0.53 | **<.0001** | **.0002** | 2.05 | 0.65 | **0.002** | **0.004** | 2.16 | 0.52 | **0.0002** | **0.001** |
| GSH | 1.31 | 1.26 | 0.152 | 0.150 | 5.61 | 1.26 | **<.0001** | **<.0001** | 3.54 | 1.35 | **0.007** | **0.008** | 4.72 | 1.08 | **<.0001** | **0.0001** |
| Glu | 0.75 | 0.60 | 0.111 | 0.220 | 2.72 | 0.64 | **0.0001** | **0.001** | 2.00 | 0.66 | **0.003** | **0.010** | 2.26 | 0.80 | **0.004** | **0.012** |
| Ins | 0.74 | 0.37 | **0.029** | **0.029** | 2.09 | 0.55 | **0.0004** | **0.001** | 1.67 | 0.54 | **0.002** | **0.004** | 2.86 | 0.70 | **0.0002** | **0.001** |
| Lac | 6.83 | 3.31 | **0.024** | **0.034** | 26.24 | 2.54 | **<.0001** | **<.0001** | 20.44 | 4.48 | **<.0001** | **0.0002** | 13.05 | 3.07 | **0.0001** | **0.0003** |
| tNAA | 0.02 | 0.20 | 0.470 | **0.047** | 0.79 | 0.28 | **0.004** | **0.012** | 0.39 | 0.30 | 0.101 | 0.160 | 0.72 | 0.29 | **0.010** | **0.019** |
| tCh | 1.73 | 0.66 | **0.007** | **0.007** | 5.13 | 0.75 | **<.0001** | **<.0001** | 3.50 | 1.12 | **0.002** | **0.002** | 3.52 | 0.95 | **0.0005** | **0.001** |
| PE | -2.15 | 1.86 | 0.129 | 0.170 | -5.16 | 1.33 | **0.0003** | **0.002** | -1.66 | 2.43 | 0.250 | 0.250 | -1.84 | 2.25 | 0.210 | 0.240 |
| Scyllo | -0.42 | 2.42 | 0.432 | 0.490 | 1.77 | 2.55 | 0.246 | 0.480 | 1.97 | 2.44 | 0.213 | 0.480 | 2.47 | 2.48 | 0.164 | 0.480 |
| Glc+Tau | -2.95 | 1.01 | **0.003** | **0.014** | -3.35 | 1.01 | **0.001** | **0.010** | -1.86 | 1.28 | 0.079 | 0.130 | -2.81 | 1.27 | **0.018** | **0.048** |
| sigma | -5.49 | 1.13 | **<.0001** | **0.0002** | -6.45 | 1.91 | **0.001** | **0.003** | -1.16 | 1.83 | 0.267 | 0.267 | 3.11 | 1.85 | 0.052 | 0.060 |
| | Block 5 | | | | Block 6 | | | | Block 7 | | | | Block 8 | | | |
| | % | ±SD | p | $p_{FDR}$ | % | ±SD | p | $p_{FDR}$ | % | ±SD | p | $p_{FDR}$ | % | ±SD | p | $p_{FDR}$ |
| Asc | 5.21 | 3.40 | **0.068** | 0.280 | 3.91 | 4.81 | **0.211** | 0.290 | 1.55 | 3.92 | **0.348** | 0.350 | 3.48 | 5.04 | 0.248 | 0.290 |
| Asp | 0.06 | 2.16 | 0.489 | 0.490 | 1.41 | 1.77 | 0.216 | 0.290 | 3.08 | 1.90 | 0.058 | 0.120 | 1.70 | 1.58 | 0.146 | 0.120 |
| tCr | 1.88 | 0.58 | **0.001** | **0.004** | 1.62 | 0.53 | **0.003** | **0.004** | 1.35 | 0.53 | **0.009** | **0.010** | 1.33 | 0.52 | **0.009** | **0.010** |
| GSH | 5.68 | 1.17 | **<.0001** | **0.0001** | 5.46 | 1.09 | **<.0001** | **0.0001** | 5.29 | 1.17 | **<.0001** | **0.0001** | 4.25 | 1.35 | **0.002** | **0.003** |
| Glu | 0.66 | 0.83 | 0.214 | 0.280 | 0.48 | 0.67 | 0.241 | 0.280 | -0.07 | 0.82 | 0.466 | 0.460 | -0.74 | 0.85 | 0.195 | 0.280 |
| Ins | 2.12 | 0.77 | **0.005** | **0.007** | 2.11 | 0.55 | **0.0004** | **0.001** | 1.67 | 0.54 | **0.002** | **0.028** | 1.24 | 0.60 | **0.025** | 0.100 |
| Lac | 11.92 | 3.79 | **0.002** | **0.004** | 8.53 | 4.19 | **0.026** | **0.034** | 2.75 | 3.08 | 0.190 | **0.019** | 4.46 | 3.26 | 0.091 | 0.100 |
| tNAA | 0.74 | 0.26 | **0.004** | **0.012** | 0.67 | 0.19 | **0.001** | **0.007** | 0.15 | 0.29 | 0.309 | **0.035** | 0.17 | 0.22 | 0.228 | **0.030** |

| | | | | | | | | | | | | | | | | |
|---|---|---|---|---|---|---|---|---|---|---|---|---|---|---|---|---|
| tCh | 4.03 | 0.95 | **0.0001** | **0.0002** | 3.70 | 0.87 | **0.0001** | **0.0002** | 3.70 | 0.97 | **0.0004** | **0.001** | 3.73 | 0.88 | **0.0001** | **0.0002** |
| PE | -4.75 | 2.15 | **0.018** | **0.042** | -4.48 | 2.21 | **0.026** | **0.042** | -5.60 | 2.67 | **0.023** | **0.042** | -4.18 | 1.91 | **0.019** | **0.042** |
| Scyllo | -0.99 | 2.74 | 0.360 | 0.480 | 0.98 | 2.73 | 0.361 | 0.480 | -1.41 | 2.52 | 0.290 | 0.480 | -0.10 | 2.63 | 0.485 | 0.490 |
| Glc+Tau | -2.69 | 1.47 | **0.039** | 0.078 | -1.56 | 1.61 | 0.170 | 0.227 | 0.04 | 1.26 | 0.489 | 0.489 | 0.47 | 1.37 | 0.368 | 0.420 |
| sigma | 6.71 | 2.03 | **0.001** | **0.003** | 7.04 | 1.94 | **0.001** | **0.002** | 6.85 | 2.30 | **0.003** | **0.004** | 6.59 | 2.15 | **0.002** | **0.004** |

Asc: ascorbate; Asp: aspartate; Glc+Tau: glucose + composite taurine; Glu: glutamate; GSH: glutathione; Ins: inositol; Lac: lactate; PE: phosphoethanolamine; tCh: total choline; tCr: total creatine; tNAA: total N-acetylaspartate.

**Table S14. phMRS: Dose-dependent effects of S-ketamine on neurometabolite levels in the ACC.** Per comparison between conditions, and per infusion block, the % change with SD are provided, as well as the original p-value and the false discovery rate (FDR)-corrected p-values ($p_{FDR}$).

ACC: anterior cingulate cortex; phMRS: pharmacological magnetic resonance spectroscopy.

**Placebo vs low dose**

| | Block 1 | | | | Block 2 | | | | Block 3 | | | | Block 4 | | | |
|---|---|---|---|---|---|---|---|---|---|---|---|---|---|---|---|---|
| | % | ±SD | p | $p_{FDR}$ | % | ±SD | p | $p_{FDR}$ | % | ±SD | p | $p_{FDR}$ | % | ±SD | p | $p_{FDR}$ |
| Asc | 7.62 | 13.15 | 0.28 | 0.405 | -11.01 | 10.47 | 0.15 | 0.405 | 5.36 | 12.57 | 0.34 | 0.405 | -9.01 | 11.32 | 0.22 | 0.405 |
| Asp | 80.55 | 62.70 | 0.11 | 0.494 | -61.03 | 55.09 | 0.14 | 0.494 | 1.00 | 66.65 | 0.49 | 0.494 | 7.26 | 61.52 | 0.45 | 0.494 |
| tCr | 8.10 | 6.95 | 0.13 | 0.128 | 16.24 | 8.07 | **0.03** | 0.449 | 24.67 | 10.02 | **0.01** | **0.029** | 18.83 | 10.68 | **0.05** | 0.061 |
| GSH | 14.84 | 3.33 | 0.50 | 0.497 | 2841.7 | 1794.3 | 0.06 | 0.352 | 1544.4 | 1888.6 | 0.21 | 0.352 | 2106.1 | 1930.6 | 0.14 | 0.352 |
| Glu | 22.02 | 13.80 | 0.06 | 0.099 | 18.40 | 17.57 | 0.15 | 0.153 | 39.66 | 15.21 | **0.01** | 0.063 | 21.87 | 15.98 | 0.09 | 0.123 |
| Ins | -0.96 | 10.16 | 0.46 | 0.463 | 8.47 | 10.80 | 0.22 | 0.294 | 12.87 | 12.83 | 0.16 | 0.261 | 13.04 | 12.29 | 0.15 | 0.261 |
| Lac | -4.39 | 37.57 | 0.45 | 0.557 | 50.22 | 34.35 | 0.08 | 0.557 | 8.21 | 36.39 | 0.41 | 0.557 | 20.34 | 38.47 | 0.30 | 0.557 |
| tNAA | 7.52 | 5.65 | 0.10 | 0.098 | 16.24 | 7.32 | **0.02** | 0.074 | 11.71 | 7.67 | 0.07 | 0.095 | 17.88 | 10.21 | **0.05** | 0.093 |
| tCh | -1.16 | 8.35 | 0.45 | 0.454 | -2.36 | 5.09 | 0.32 | 0.432 | 7.62 | 7.65 | 0.16 | 0.432 | 1.44 | 8.99 | 0.44 | 0.445 |
| PE | -7.84 | 19.93 | 0.35 | 0.399 | -36.48 | 13.29 | **0.01** | **0.023** | -16.67 | 15.25 | 0.14 | 0.190 | -49.94 | 16.81 | **0.01** | **0.023** |
| Scyllo | 46.80 | 44.83 | 0.15 | 0.439 | 15.83 | 45.04 | 0.36 | 0.439 | -11.28 | 45.23 | 0.40 | 0.439 | 70.45 | 45.35 | 0.07 | 0.439 |
| Glc+Tau | -21.18 | 22.93 | 0.18 | 0.395 | 17.07 | 30.70 | 0.29 | 0.395 | -8.85 | 27.05 | 0.37 | 0.395 | 12.50 | 23.56 | 0.30 | 0.395 |
| sigma | -21.64 | 7.93 | **0.01** | **0.048** | -30.76 | 12.93 | **0.01** | 0.052 | -15.87 | 15.33 | 0.16 | 0.178 | -18.77 | 12.95 | 0.08 | 0.107 |

| | Block 5 | | | | Block 6 | | | | Block 7 | | | | Block 8 | | | |
|---|---|---|---|---|---|---|---|---|---|---|---|---|---|---|---|---|
| | % | ±SD | p | $p_{FDR}$ | % | ±SD | p | $p_{FDR}$ | % | ±SD | p | $p_{FDR}$ | % | ±SD | p | $p_{FDR}$ |
| Asc | 4.62 | 12.21 | 0.35 | 0.405 | -24.59 | 13.65 | 0.339 | 0.34 | -14.17 | 11.99 | 0.12 | 0.405 | 1.63 | 12.89 | 0.45 | 0.450 |
| Asp | 17.57 | 68.56 | 0.40 | 0.494 | -1.76 | 65.93 | 0.494 | 0.49 | -49.97 | 66.59 | 0.23 | 0.494 | 33.26 | 66.46 | 0.31 | 0.494 |
| tCr | 32.12 | 9.72 | **0.002** | **0.012** | 22.49 | 8.42 | **0.027** | **0.03** | 18.54 | 11.18 | 0.06 | 0.063 | 25.13 | 11.20 | **0.02** | **0.035** |
| GSH | 1817.3 | 1633.6 | 0.14 | 0.352 | -1567.7 | 1995.1 | 0.352 | 0.35 | 803.44 | 1991.5 | 0.35 | 0.395 | 990.95 | 1728.6 | 0.29 | 0.381 |
| Glu | 39.47 | 20.45 | **0.03** | 0.079 | 31.48 | 13.94 | 0.068 | 0.07 | 32.30 | 17.54 | **0.04** | 0.079 | 22.92 | 19.95 | 0.13 | 0.150 |
| Ins | 20.42 | 14.82 | 0.09 | 0.261 | 4.68 | 13.91 | 0.423 | 0.42 | 17.76 | 11.57 | 0.07 | 0.261 | 16.20 | 15.11 | 0.15 | 0.261 |
| Lac | 21.03 | 39.51 | 0.30 | 0.557 | -26.61 | 35.03 | 0.772 | 0.77 | 14.58 | 37.16 | 0.35 | 0.557 | 1.55 | 49.92 | 0.49 | 0.557 |
| tNAA | 23.15 | 9.25 | **0.01** | 0.074 | 18.44 | 9.48 | 0.085 | 0.09 | 15.54 | 10.86 | 0.08 | 0.095 | 17.34 | 11.46 | 0.07 | 0.095 |
| tCh | 19.50 | 7.73 | **0.01** | 0.076 | 10.95 | 7.64 | 0.331 | 0.33 | 5.18 | 9.86 | 0.30 | 0.432 | 6.60 | 9.28 | 0.24 | 0.432 |

| | | | | | | | | | | | | | | | | |
|---|---|---|---|---|---|---|---|---|---|---|---|---|---|---|---|---|
| PE | 0.14 | 18.44 | 0.50 | 0.497 | -20.98 | 16.06 | 0.164 | 0.16 | -35.60 | 19.05 | **0.04** | 0.075 | -31.31 | 15.11 | **0.02** | 0.066 |
| Scyllo | -10.05 | 45.43 | 0.41 | 0.439 | 30.18 | 50.73 | 0.439 | 0.44 | 7.12 | 45.78 | 0.44 | 0.439 | 10.76 | 51.97 | 0.42 | 0.439 |
| Glc+Tau | 7.78 | 28.81 | 0.39 | 0.395 | 16.74 | 25.19 | 0.395 | 0.39 | 43.03 | 26.41 | 0.06 | 0.395 | 10.31 | 29.58 | 0.37 | 0.395 |
| sigma | -7.81 | 11.80 | 0.26 | 0.257 | -20.90 | 12.78 | 0.06 | 0.093 | -33.79 | 17.01 | **0.03** | 0.061 | -37.32 | 18.90 | **0.03** | 0.061 |

**Placebo vs high dose**

| | Block 1 | | | | Block 2 | | | | Block 3 | | | | Block 4 | | | |
|---|---|---|---|---|---|---|---|---|---|---|---|---|---|---|---|---|
| | % | ±SD | p | $p_{FDR}$ | % | ±SD | p | $p_{FDR}$ | % | ±SD | p | $p_{FDR}$ | % | ±SD | p | $p_{FDR}$ |
| Asc | 22.69 | 23.81 | 0.18 | 0.38 | -18.15 | 27.28 | 0.256 | 0.382 | 15.79 | 24.15 | 0.26 | 0.38 | 12.43 | 30.98 | 0.346 | 0.395 |
| Asp | -12.39 | 10.04 | 0.12 | 0.44 | -18.06 | 11.62 | 0.067 | 0.438 | -10.97 | 11.44 | 0.17 | 0.44 | -6.70 | 10.00 | 0.255 | 0.438 |
| tCr | 14.01 | 6.50 | **0.02** | **0.02** | 28.18 | 8.73 | **0.002** | **0.004** | 26.92 | 11.42 | **0.01** | **0.02** | 34.63 | 11.72 | **0.004** | **0.005** |
| GSH | 6.14 | 15.68 | 0.35 | 0.41 | 17.96 | 15.19 | 0.124 | 0.249 | 1.55 | 21.17 | 0.47 | 0.47 | 5.58 | 15.39 | 0.360 | 0.442 |
| Glu | 17.54 | 12.61 | 0.09 | 0.09 | 24.03 | 13.51 | **0.044** | 0.051 | 38.46 | 14.31 | **0.01** | **0.01** | 43.36 | 14.95 | **0.004** | **0.013** |
| Ins | 7.67 | 10.64 | 0.24 | 0.24 | 26.05 | 12.84 | **0.027** | **0.039** | 30.10 | 15.73 | **0.03** | **0.04** | 46.59 | 15.72 | **0.003** | **0.028** |
| Lac | 11.87 | 15.45 | 0.23 | 0.23 | 99.22 | 16.59 | **<.0001** | **<.0001** | 49.48 | 24.53 | **0.03** | 0.08 | 43.50 | 22.27 | **0.032** | 0.083 |
| tNAA | 4.40 | 8.73 | 0.31 | 0.31 | 27.62 | 9.60 | **0.004** | **0.007** | 12.99 | 12.46 | 0.15 | 0.18 | 37.08 | 13.59 | **0.006** | **0.008** |
| tCh | 12.98 | 10.87 | 0.12 | 0.12 | 50.92 | 12.93 | **0.0003** | **0.003** | 29.06 | 15.05 | **0.03** | **0.04** | 35.37 | 15.53 | **0.016** | **0.022** |
| PE | 10.32 | 37.33 | 0.39 | 0.45 | 31.20 | 27.34 | 0.133 | 0.338 | -31.35 | 27.41 | 0.13 | 0.34 | -34.26 | 36.52 | 0.179 | 0.338 |
| Scyllo | 14.61 | 27.20 | 0.30 | 0.33 | 32.11 | 27.42 | 0.127 | 0.243 | 43.15 | 29.68 | 0.08 | 0.21 | 93.55 | 27.60 | **0.001** | **0.010** |
| Glc+Tau | -73.20 | 47.51 | 0.07 | 0.27 | -26.42 | 57.34 | 0.325 | 0.389 | -27.15 | 65.08 | 0.34 | 0.39 | -47.67 | 58.71 | 0.213 | 0.340 |
| sigma | -109.4 | 24.04 | **<.0001** | **0.0006** | -181.1 | 43.92 | **0.0002** | **0.0008** | -79.48 | 54.73 | 0.08 | 0.13 | -5.40 | 51.45 | 0.459 | 0.459 |
| | Block 5 | | | | Block 6 | | | | Block 7 | | | | Block 8 | | | |
| | % | ±SD | p | $p_{FDR}$ | % | ±SD | p | $p_{FDR}$ | % | ±SD | p | $p_{FDR}$ | % | ±SD | p | $p_{FDR}$ |
| Asc | 17.95 | 29.54 | 0.275 | 0.382 | -32.44 | 24.19 | 0.096 | 0.382 | -17.77 | 31.05 | 0.286 | 0.382 | -1.98 | 31.25 | 0.475 | 0.475 |
| Asp | -4.31 | 14.37 | 0.383 | 0.438 | -0.36 | 10.02 | 0.486 | 0.486 | 4.87 | 12.84 | 0.354 | 0.438 | 4.11 | 10.04 | 0.343 | 0.438 |
| tCr | 36.38 | 11.01 | **0.002** | **0.004** | 32.80 | 10.52 | **0.002** | **0.004** | 30.45 | 9.52 | **0.002** | **0.004** | 41.23 | 10.43 | **0.000** | **0.003** |
| GSH | 18.75 | 15.43 | 0.118 | 0.249 | 35.70 | 18.04 | **0.030** | 0.177 | 32.69 | 18.39 | **0.044** | 0.177 | 9.62 | 15.49 | 0.270 | 0.412 |
| Glu | 40.54 | 16.07 | **0.010** | **0.015** | 37.66 | 11.85 | **0.002** | **0.013** | 38.25 | 13.64 | **0.005** | **0.013** | 34.78 | 14.31 | **0.012** | **0.016** |
| Ins | 37.36 | 18.78 | **0.029** | **0.039** | 31.82 | 14.63 | **0.020** | **0.039** | 31.44 | 12.38 | **0.009** | **0.037** | 31.15 | 16.17 | **0.033** | **0.039** |
| Lac | 43.06 | 23.75 | **0.041** | 0.083 | 21.94 | 22.00 | 0.164 | 0.230 | 18.94 | 19.62 | 0.172 | 0.230 | 18.38 | 24.75 | 0.233 | 0.233 |
| tNAA | 38.60 | 13.13 | **0.004** | **0.007** | 35.83 | 12.44 | **0.004** | **0.007** | 42.74 | 15.04 | **0.005** | **0.007** | 43.69 | 12.65 | **0.001** | **0.007** |
| tCh | 50.22 | 16.62 | **0.003** | **0.012** | 37.88 | 14.00 | **0.006** | **0.016** | 35.03 | 13.52 | **0.008** | **0.016** | 38.03 | 15.40 | **0.011** | **0.017** |

| | | | | | | | | | | | | | | | | |
|---|---|---|---|---|---|---|---|---|---|---|---|---|---|---|---|---|
| PE | 18.04 | 36.10 | 0.311 | 0.415 | -1.47 | 39.15 | 0.485 | 0.485 | -27.04 | 33.08 | 0.211 | 0.338 | -67.45 | 32.59 | **0.025** | 0.200 |
| Scyllo | 19.10 | 27.66 | 0.248 | 0.330 | 30.99 | 29.48 | 0.152 | 0.243 | 12.34 | 27.72 | 0.330 | 0.330 | 60.74 | 27.74 | **0.019** | 0.078 |
| Glc+Tau | -60.39 | 60.21 | 0.163 | 0.326 | 9.24 | 72.33 | 0.450 | 0.447 | 106.59 | 69.21 | 0.069 | 0.274 | 60.15 | 57.86 | 0.155 | 0.326 |
| sigma | 73.07 | 54.44 | 0.096 | 0.128 | 75.92 | 53.87 | 0.086 | 0.128 | 87.36 | 65.00 | 0.096 | 0.144 | 77.33 | 65.68 | 0.126 | 0.144 |

**Low dose vs high dose**

| | Block 1 | | | | Block 2 | | | | Block 3 | | | | Block 4 | | | |
|---|---|---|---|---|---|---|---|---|---|---|---|---|---|---|---|---|
| | % | ±SD | p | $p_{FDR}$ | % | ±SD | p | $p_{FDR}$ | % | ±SD | p | $p_{FDR}$ | % | ±SD | p | $p_{FDR}$ |
| Asc | -5.17 | 18.74 | 0.393 | 0.480 | -4.80 | 17.84 | 0.395 | 0.480 | -3.46 | 20.37 | 0.433 | 0.480 | -21.49 | 19.46 | 0.140 | 0.480 |
| Asp | -42.20 | 14.86 | **0.005** | **0.037** | -12.62 | 15.41 | 0.211 | 0.281 | -20.72 | 16.07 | 0.105 | 0.281 | -17.00 | 15.00 | 0.134 | 0.281 |
| tCr | 29.16 | 21.94 | 0.098 | 0.118 | 53.02 | 26.81 | **0.030** | 0.080 | 27.49 | 37.02 | 0.233 | 0.233 | 72.50 | 31.50 | **0.015** | 0.076 |
| GSH | -1.96 | 15.61 | 0.451 | 0.451 | 45.85 | 19.18 | **0.013** | **0.051** | 17.47 | 18.98 | 0.183 | 0.245 | 29.74 | 18.39 | 0.060 | 0.106 |
| Glu | 12.97 | 49.36 | 0.398 | 0.398 | 33.60 | 72.48 | 0.324 | 0.398 | 23.48 | 67.92 | 0.366 | 0.398 | 118.83 | 65.33 | **0.041** | 0.328 |
| Ins | 67.56 | 98.01 | 0.249 | 0.249 | 126.83 | 135.0 | 0.179 | 0.204 | 148.49 | 126.6 | 0.126 | 0.204 | 295.94 | 137.8 | **0.021** | 0.170 |
| Lac | -10.04 | 90.85 | 0.456 | 0.457 | 249.94 | 78.32 | **0.002** | **0.016** | 140.36 | 104.7 | 0.097 | 0.193 | 131.70 | 69.83 | **0.036** | 0.144 |
| tNAA | 30.00 | 38.91 | 0.224 | 0.299 | -20.66 | 58.87 | 0.364 | 0.417 | 6.05 | 48.71 | 0.451 | 0.451 | -59.02 | 52.77 | 0.137 | 0.272 |
| tCh | -24.14 | 16.91 | 0.083 | 0.095 | -80.52 | 17.66 | **<.0001** | **0.001** | -29.27 | 27.53 | 0.149 | 0.149 | -57.05 | 23.85 | **0.013** | 0.051 |
| PE | -13.95 | 24.40 | 0.286 | 0.417 | -66.61 | 20.43 | **0.002** | **0.014** | -6.27 | 21.71 | 0.388 | 0.443 | -43.63 | 26.31 | 0.055 | 0.222 |
| Scyllo | 3.82 | 110.8 | 0.486 | 0.486 | 126.40 | 123.7 | 0.159 | 0.311 | 238.74 | 120.2 | **0.030** | 0.140 | 176.38 | 112.4 | 0.065 | 0.174 |
| Glc+Tau | 30.33 | 33.82 | 0.190 | 0.309 | 49.50 | 34.59 | 0.083 | 0.221 | 25.66 | 34.44 | 0.232 | 0.309 | 65.29 | 34.03 | **0.034** | 0.221 |
| sigma | 28.18 | 13.84 | **0.027** | **0.043** | 54.59 | 22.50 | **0.012** | **0.030** | 27.25 | 21.55 | 0.109 | 0.125 | -16.50 | 20.50 | 0.215 | 0.215 |

| | Block 5 | | | | Block 6 | | | | Block 7 | | | | Block 8 | | | |
|---|---|---|---|---|---|---|---|---|---|---|---|---|---|---|---|---|
| | % | ±SD | p | $p_{FDR}$ | % | ±SD | p | $p_{FDR}$ | % | ±SD | p | $p_{FDR}$ | % | ±SD | p | $p_{FDR}$ |
| Asc | -3.77 | 17.45 | 0.416 | 0.480 | -23.76 | 23.76 | 0.164 | 0.480 | -11.17 | 21.75 | 0.306 | 0.480 | 1.22 | 24.20 | 0.480 | 0.480 |
| Asp | -17.30 | 19.77 | 0.195 | 0.281 | -7.96 | 15.52 | 0.306 | 0.314 | 15.21 | 17.20 | 0.193 | 0.281 | -9.35 | 19.03 | 0.314 | 0.314 |
| tCr | 40.68 | 31.25 | 0.103 | 0.118 | 47.39 | 27.89 | 0.051 | 0.103 | 47.74 | 33.46 | 0.084 | 0.177 | 70.39 | 32.00 | **0.019** | 0.076 |
| GSH | 38.61 | 18.85 | **0.026** | 0.070 | 13.94 | 21.22 | 0.259 | 0.296 | 42.59 | 16.73 | **0.009** | 0.051 | 25.66 | 16.46 | 0.066 | 0.106 |
| Glu | 33.54 | 58.97 | 0.288 | 0.398 | 40.36 | 56.24 | 0.240 | 0.398 | 31.03 | 66.52 | 0.323 | 0.398 | 63.41 | 67.45 | 0.178 | 0.398 |
| Ins | 161.45 | 123.9 | 0.103 | 0.204 | 225.22 | 130.4 | **0.049** | 0.195 | 117.46 | 124.5 | 0.178 | 0.204 | 150.13 | 142.2 | 0.151 | 0.204 |
| Lac | 107.00 | 71.37 | 0.074 | 0.193 | 118.47 | 104.4 | 0.134 | 0.215 | 11.67 | 82.66 | 0.444 | 0.457 | 28.88 | 79.15 | 0.359 | 0.457 |
| tNAA | -44.08 | 45.20 | 0.170 | 0.272 | -58.44 | 44.73 | 0.102 | 0.272 | -97.41 | 61.07 | 0.062 | 0.249 | -96.28 | 52.69 | **0.040** | 0.249 |
| tCh | -36.83 | 24.13 | 0.070 | 0.095 | -38.00 | 21.78 | **0.047** | 0.095 | -39.61 | 27.58 | 0.082 | 0.095 | -44.08 | 26.39 | 0.054 | 0.095 |

| | | | | | | | | | | | | | | | | |
|---|---|---|---|---|---|---|---|---|---|---|---|---|---|---|---|---|
| PE | -13.66 | 27.56 | 0.312 | 0.417 | -24.20 | 21.41 | 0.135 | 0.360 | -24.37 | 26.28 | 0.182 | 0.363 | 1.58 | 24.37 | 0.474 | 0.475 |
| Scyllo | 99.17 | 113.0 | 0.195 | 0.311 | 71.73 | 134.2 | 0.299 | 0.395 | 54.29 | 134.7 | 0.345 | 0.395 | 229.75 | 120.8 | **0.035** | 0.140 |
| Glc+Tau | 63.04 | 40.24 | 0.065 | 0.212 | 28.65 | 37.46 | 0.226 | 0.309 | -6.89 | 44.94 | 0.440 | 0.440 | -19.73 | 45.02 | 0.333 | 0.380 |
| sigma | -43.20 | 23.37 | **0.039** | 0.052 | -63.20 | 27.30 | **0.015** | **0.030** | -87.41 | 34.39 | **0.009** | **0.030** | -88.86 | 31.60 | **0.005** | **0.030** |

Asc: ascorbate; Asp: aspartate; Glc+Tau: glucose + composite taurine; Glu: glutamate; GSH: glutathione; Ins: inositol; Lac: lactate; PE: phosphoethanolamine; tCh: total choline; tCr: total creatine; tNAA: total N-acetylaspartate.

**Table S15. Bayesian multilevel modeling to estimate condition based on individual-level phMRI contrast of parameter estimates in the ACC. A.** Coefficients for the best-fitting Bayesian model with categorical outcome variable (condition) and phMRI COPE as predictor. Leave-one-out information criterion (LOO-IC): 1119.5±19.0; widely applicable information criterion (WAIC): 1119.5±19.0. B. Differences in predictor estimates extracted from the posterior distributions for low and high S-ketamine dose conditions. For all measures, the estimate and 95% confidence intervals are provided. Model: condition ~ COPE_fMRI + (1 | Subject)

ACC: anterior cingulate cortex; COPE: contrast of parameter estimate; phMRI: pharmacological magnetic resonance imaging.

| **A. Model estimates for low and high dose compared to placebo** | | |
|---|---|---|
| | **Placebo vs low dose** | **Placebo vs high dose** |
| Multilevel hyperparameters | | |
| intercept | **Est: 0.09; CI: 0.00-0.25** | **Est: 0.10; CI: 0.00-0.30** |
| Regression coefficients | | |
| intercept | Est: -0.15; CI: -0.37-0.08 | **Est: -0.26; CI: -0.49 - -0.02** |
| phMRI | **Est: 0.03; CI: 0.02-0.04** | **Est: 0.04; CI: 0.03-0.05** |

| **B. Difference scores for high dose compared to low dose** | |
|---|---|
| | **Low vs high dose** |
| Regression coefficients | |
| phMRI | Est: 0.005; CI: -0.0007-0.011 |

**Table S16. Bayesian multilevel modeling to estimate condition based on individual-level phMRS contrast of parameter estimates in the ACC. A.** Coefficients for the best-fitting Bayesian model with categorical outcome variable (condition) and glutamate (Glu), lactate (Lac), aspartate (Asp), and glucose (Glc) COPE as predictors. LOO-IC: 1182.4±; WAIC: 182.3±15.5. **B.** Differences in predictor estimates extracted from the posterior distributions for low and high S-ketamine dose conditions. For all measures, the estimate and 95% confidence intervals are provided.

Model: condition ~ COPE_Glu + COPE_Asp + COPE_Lac + COPE_Glc + (1 | Subject)

ACC: anterior cingulate cortex; COPE: contrast of parameter estimate; phMRS: pharmacological magnetic resonance spectroscopy.

| **A. Model estimates for low and high dose compared to placebo** | | |
|---|---|---|
| | **Placebo vs low dose** | **Placebo vs high dose** |
| Multilevel hyperparameters | | |
| intercept | **Est: 0.09; CI: 0.00-0.26** | **Est: 0.10; CI: 0.00-0.29** |
| Regression coefficients | | |
| intercept | Est: 0.10; CI: -0.13-0.34 | Est: -0.10; CI: -0.35-0.15 |
| Glu | **Est: 41.63; CI: 24.26-59.59** | **Est: 41.35; CI: 23.73-59.77** |
| Lac | Est: -24.76; CI: -63.50-13.84 | **Est: 42.60; CI: 2.99-81.66** |
| Asp | Est: 1.01; CI: -12.30-14.58 | Est: -12.07; CI: -25.77-1.35 |
| Glc | Est: 23.27; CI: -7.03-53.04 | Est: 2.42; CI: -26.89-31.98 |
| **B. Difference scores for high dose compared to low dose** | | |
| **Low vs high dose** | | |
| Regression coefficients | | |
| Glu | Est: -0.27; CI: -18.39-17.74 | |
| Lac | Est: 67.36; CI: 30.00-105.44 | |
| Asp | Est: -13.08; CI: -26.70-0.38 | |
| Glc | Est: -20.86; CI: -49.88-7.12 | |

Asp: aspartate; Glc: glucose + composite taurine; Glu: glutamate; Lac: lactate.

**Table S17. Bayesian multilevel modeling to estimate condition based on individual-level combined phMRI+phMRS contrast of parameter estimates in the ACC. A.** Coefficients for the best-fitting Bayesian model with categorical outcome variable (condition) and phMRI, glutamate (Glu), and lactate (Lac) COPE as predictors. LOO-IC: 1082.2±23.0; WAIC: 1082.1±23.0. **B.** Differences in predictor estimates extracted from the posterior distributions for low and high S-ketamine dose conditions. . For all measures, the estimate and 95% confidence intervals are provided.
Model: condition ~ COPE_fMRI + COPE_Glu + COPE_Asp + COPE_Lac + COPE_Glc + (1 | Subject)

ACC: anterior cingulate cortex; COPE: contrast of parameter estimate; phMRI: pharmacological magnetic resonance imaging; phMRS: pharmacological magnetic resonance spectroscopy

| **A. Model estimates for low and high dose compared to placebo** | | |
|---|---|---|
| | **Placebo vs low dose** | **Placebo vs high dose** |
| Multilevel hyperparameters | | |
| intercept | **Est: 0.09; CI: 0.00-0.29** | **Est: 0.14; CI: 0.01-0.40** |
| Regression coefficients | | |
| intercept | Est: -0.10; CI: -0.34-0.15 | **Est: -0.41; CI: -0.67- -0.15** |
| phMRI | **Est: 0.03; CI: 0.02-0.04** | **Est: 0.04; CI: 0.03-0.05** |
| Glu | **Est: 39.40; CI: 21.35-58.05** | **Est: 43.60; CI: 24.04-62.92** |
| Lac | Est: -11.21; CI: -52.36-30.52 | **Est: 56.96; CI: 15.26-100.02** |

| **B. Difference scores for high dose compared to low dose** | |
|---|---|
| | **Low vs high dose** |
| Regression coefficients | |
| phMRI | **Est: 0.006; CI: 0.0002-0.012** |
| Glu | Est: 4.20; CI: -13.00-21.37 |
| Lac | **Est: 68.17; CI: 31.42-105.44** |

Glu: glutamate; Lac: lactate.

**Table S18. Association between subject-level neuroimaging measures and dissociation. A.** Per infusion block, the results from the repeated measures correlations between the phMRI COPE values and total dissociation score, measured using the CADSS, are provided. **B.** Per infusion block, results from the repeated measures correlations between phMRS COPE values and total CADSS score per metabolite.

COPE: contrast of parameter estimates; phMRI: pharmacological magnetic resonance imaging; phMRS: pharmacological magnetic resonance spectroscopy.

| **A. phMRI** | | | | | | | | | | | | |
|---|---|---|---|---|---|---|---|---|---|---|---|---|
| | Block 1 (0 - 4 min) | | | Block 2 (4 – 8 min) | | | Block 3 (8 – 12 min) | | | Block 4 (12 – 16 min) | | |
| | r | p | $p_{FDR}$ | r | p | $p_{FDR}$ | r | p | $p_{FDR}$ | r | p | $p_{FDR}$ |
| **phMRI** | .42 | **.003** | **.005** | .70 | **<.0001** | **<.0001** | .30 | **.03** | **.03** | .36 | **.01** | **.01** |
| | Block 5 (16 – 20 min) | | | Block 6 (20 – 24 min) | | | Block 7 (24 – 28 min) | | | Block 8 (28 – 32 min) | | |
| | r | p | $p_{FDR}$ | r | p | $p_{FDR}$ | r | p | $p_{FDR}$ | r | p | $p_{FDR}$ |
| **phMRI** | .37 | **.007** | **.01** | .48 | **.0004** | **.001** | .49 | **.0002** | **.001** | .37 | **.008** | **.01** |

| **B. phMRS** | | | | | | | | | | | | |
|---|---|---|---|---|---|---|---|---|---|---|---|---|
| | Block 1 (0 - 4 min) | | | Block 2 (4 – 8 min) | | | Block 3 (8 – 12 min) | | | Block 4 (12 – 16 min) | | |
| | r | p | $p_{FDR}$ | r | p | $p_{FDR}$ | r | p | $p_{FDR}$ | r | p | $p_{FDR}$ |
| **Glu** | 0.33 | **0.02** | 0.08 | 0.30 | **0.04** | 0.08 | 0.31 | **0.03** | 0.08 | 0.25 | 0.08 | 0.11 |
| **Asp** | -0.03 | 0.83 | 0.95 | -0.22 | 0.13 | 0.86 | -0.15 | 0.32 | 0.86 | -0.13 | 0.39 | 0.86 |
| **Glc** | -0.16 | 0.29 | 0.65 | -0.02 | 0.88 | 0.88 | 0.02 | 0.88 | 0.88 | 0.09 | 0.55 | 0.88 |
| **Lac** | -0.03 | 0.83 | 0.83 | -0.20 | 0.17 | 0.83 | -0.15 | 0.32 | 0.83 | -0.13 | 0.39 | 0.83 |
| | Block 5 (16 – 20 min) | | | Block 6 (20 – 24 min) | | | Block 7 (24 – 28 min) | | | Block 8 (28 – 32 min) | | |
| | r | p | $p_{FDR}$ | r | p | $p_{FDR}$ | r | p | $p_{FDR}$ | r | p | $p_{FDR}$ |
| **Glu** | 0.21 | 0.15 | 0.14 | 0.30 | **0.04** | 0.08 | 0.25 | 0.08 | 0.11 | 0.22 | 0.13 | 0.14 |
| **Asp** | 0.01 | 0.95 | 0.95 | -0.10 | 0.48 | 0.86 | -0.04 | 0.81 | 0.95 | -0.09 | 0.36 | 0.86 |
| **Glc** | 0.03 | 0.83 | 0.88 | 0.14 | 0.33 | 0.65 | 0.33 | **0.02** | 0.16 | 0.18 | 0.21 | 0.65 |
| **Lac** | -0.03 | 0.82 | 0.83 | -0.10 | 0.49 | 0.83 | -0.04 | 0.81 | 0.83 | -0.09 | 0.54 | 0.83 |

Asp: aspartate; Glc: glucose + composite taurine; Glu: glutamate; Lac: lactate.

**Table S19. Associations between phMRI outcomes and receptor distribution maps.** The t-stat maps from the low and high S-ketamine dose compared to placebo were extracted and fed into a spin-test in neuromaps, to assess the correlation of these outcomes with relevant neurotransmitter receptor maps. To assess initial and delayed responses, both infusion blocks two and five were assessed for each condition and placebo-contrast.

| **Low dose > placebo** | | | | |
|---|---|---|---|---|
| | Block 2 (8 – 12 minutes) | | Block 5 (16 – 20 minutes) | |
| | r | p | r | p |
| Glutamate MR5 | 0.40 | **0.0099** | 0.16 | 0.2277 |
| $GABA_A$ receptor | 0.0 | 1.0 | 0.20 | **0.0297** |
| NET | 0.25 | 0.1782 | 0.04 | 0.8218 |
| DAT | -0.03 | 0.7624 | -0.10 | 0.2178 |
| D2 | -0.11 | 0.5347 | -0.14 | 0.3681 |
| VAChT | 0.03 | 0.7822 | -0.11 | 0.3960 |
| mAChR M1 | 0.07 | 0.5050 | 0.04 | 0.6535 |
| 5-HTT | -0.18 | 0.2673 | -0.11 | 0.5545 |
| 5-HT2A | -0.05 | 0.5545 | 0.07 | 0.4455 |
| Opioid μ | 0.37 | 0.1980 | -0.05 | 0.7822 |
| Opioid $\kappa$ | 0.48 | **0.0198** | 0.07 | 0.7624 |
| **High dose > placebo** | | | | |
| | Block 2 (8 – 12 minutes) | | Block 5 (16 – 20 minutes) | |
| | r | p | r | p |
| Glutamate MR5 | 0.40 | **0.0099** | 0.26 | **0.0099** |
| $GABA_A$ receptor | -0.03 | 0.7822 | 0.10 | 0.3069 |
| NET | 0.23 | 0.1980 | 0.07 | 0.7129 |
| DAT | -0.02 | 0.9307 | -0.05 | 0.6436 |
| Dopamine D2 | -0.08 | 0.6733 | -0.09 | 0.5911 |
| VAChT | 0.07 | 0.5545 | -0.02 | 0.8317 |
| mAChR M1 | 0.02 | 0.8218 | 0.01 | 0.8614 |
| SERT | -0.14 | 0.4752 | -0.10 | 0.3069 |
| SER2A | -0.02 | 0.7129 | 0.03 | 0.7030 |
| Opioid μ | 0.47 | **0.0396** | 0.19 | 0.4752 |
| Opioid $\kappa$ | 0.52 | **0.0099** | 0.22 | 0.2673 |
| **High dose > low dose** | | | | |
| | Block 2 (8 – 12 minutes) | | Block 5 (16 – 20 minutes) | |
| | r | p | r | p |
| Glu MR5 | 0.31 | **0.0099** | 0.29 | **0.0099** |
| $GABA_A$ | -0.10 | 0.3069 | 0.04 | 0.7624 |
| NET | 0.15 | 0.4752 | 0.13 | 0.5446 |
| DAT | -0.05 | 0.6436 | -0.04 | 0.7327 |
| Dopamine D2 | -0.07 | 0.6832 | -0.02 | 0.9208 |
| VAChT | 0.03 | 0.8119 | 0.0 | 1.0 |

| | | | | |
|---|---|---|---|---|
| mAChR M1 | 0.06 | 0.6139 | 0.09 | 0.3267 |
| SERT | -0.11 | 0.5545 | -0.06 | 0.7525 |
| SER2A | 0.02 | 0.7228 | 0.09 | 0.3069 |
| Opioid μ | 0.39 | 0.1683 | 0.20 | 0.4752 |
| Opioid $\kappa$ | 0.35 | 0.0792 | 0.21 | 0.3069 |

Glu MR5: glutamate metabotropic receptor 5; 5-HTT: serotonin transporter; 5-HT2A: serotonin 2A receptor; VAChT: acetylcholine transporter; mAChR M1: metabotropic acetylcholine receptor 1; DAT: dopamine transporter; D2: dopamine D2 receptor, NET: norepinephrine transporter.

**Table S20. phMRI and phMRS data quality metrics.** Per condition, the full-width at half maximum (FWHM; in Hz) of total creatine (tCr), the Cramér-Rao lower bounds (CRLB) of glutamate (Glu), and the mean framewise displacement (FD; in mm) after scrubbing are provided as well as results from a repeated measures ANOVA to determine dose-dependent differences in these quality metrics. ANOVA: analysis of variance; phMRI: pharmacological magnetic resonance imaging; phMRS: pharmacological magnetic resonance spectroscopy.

| | **Placebo** | **Low dose** | **High dose** | Statistics | |
|---|---|---|---|---|---|
| | | | | *F(df)* | *p* |
| $FWHM_{tCr}$ | 13.36 (2.2) | 14.05 (2.0) | 13.59 (1.51) | F(2,50) = 0.36 | 0.70 |
| $CRLB_{Glu}$ | 2.77 (0.26) | 2.80 (0.31) | 2.90 (0.32) | F(2,50) = 1.33 | 0.57 |
| Mean FD | 0.21 (0.07) | 0.21 (0.06) | 0.23 (0.07) | F(2,50) = 0.34 | 0.71 |

**Table S21. MRSinMRS acquisition parameters.**

| | sLASER |
|---|---|
| **1. Hardware** | |
| a. Field Strenght | 7T |
| b. Manufacturer | Philips |
| c. Model | Achieva |
| d. RF coil | Quadrature birdcage transmit and 32-channel phased-array receive coils (Nova Medical) |
| e. Additional hardware | N/A |
| **2. Acquisition** | |
| a. Pulse sequence | sLASER (Interleaved with) 3D-EPI |
| b. Volume of interest | dACC |
| c. Nominal VOI size | 30 x 20 x 15 mm |
| d. Repitition time (TR), echo time (TE) | TR/TE=3500/36 ms |
| e. Total number of acquisitions/averages | 448 |
| f. Additional sequence parameters | 3000 Hz; 1024 data points |
| g. Water supression method | VAPOR |
| h. Shimming method | HOS-DLT (Boer et al. 2020) |
| i. Triggering or motion correction method | None |
| **3. Data analysis methods and outputs** | |
| a. Analysis software | FSL-MRS v.2.1.13 |
| b. Processing steps deviating from quoted product | None |
| c. Output measure | Dynamic fitting on unscaled spectra |
| d. Quantification references and assumptions | Basisset generated for this sequence (Schrantee et al. 2023) |
| 4. **Data quality** | |
| a. Reported variables | FWHM, CRLB |
| b. Data exclusion criteria | Metabolites not evaluated when >50% subjects >20% CRLB; Spectra were excluded when CRLB Glu>5% or tCr FWHM >19Hz. |
| c Quality measures of postprocessing model fitting | Glu CRLB, tCr FWHM |
| d. Sample spectrum | Figure 3A |
